# Long Terminal Repeats of Gammaretroviruses Retain Stable Expression After Integration Retargeting or Knock-In into the Restrictive Chromatin of Lamina-Associated Domains

**DOI:** 10.1101/2024.05.30.596639

**Authors:** Dalibor Miklík, Martina Slavková, Dana Kučerová, Chahrazed Mekadim, Jakub Mrázek, Jiří Hejnar

**Author notes:** The first two authors contributed equally to this work.

## Abstract

Retroviruses integrate their genomes into the genomes of infected host cells and form a genetic platform for stable gene expression. Epigenetic silencing can, however, hamper the expression of integrated provirus. As gammaretroviruses (γRVs) preferentially integrate into sites of active promoters and enhancers, the high expression activity of γRVs can be attributed to the integration preference. Long terminal repeats (LTRs) of some γRVs were shown to act as potent promoters for gene expression. Here, we investigate the capacity of different γRV LTRs to drive stable expression inside a non-preferred epigenomic environment using diverse retroviral vectors and CRISPR-Cas9-directed vector knock-in. We demonstrate that different γRV LTRs are either rapidly silenced or long-term active with active proviral population prevailing under normal and retargeted integration. In addition, we show that lamina-associated domains (LADs) can be targeted by CRISPR-Cas9 for vector insertion leading to γRV LTR-driven long-term stable gene expression. Alternatively to established γRV systems, the LTRs of feline leukemia virus and koala retrovirus are capable of driving stable, albeit intensity-diverse, transgene expression in LADs. Altogether, we show that despite the occurrence of rapid silencing events, the majority of γRV LTRs can drive stable expression after retrovirus integration or CRISPR-Cas9-directed knock-in outside of the preferred chromatin landscape.

## INTRODUCTION

Retroviruses are ssRNA viruses of the *Retroviridae* family that utilize reverse transcription and subsequent genomic DNA integration in their replication cycle. After integration, the provirus becomes a permanent part of the host genome and forms a genetic platform for host transcription machinery-driven long-term stable expression of genes encoded by the retroviral genome. The establishment of the permanent infection of the host cells makes retroviruses hard-to-cure pathogens as well as useful tools for gene delivery. The host-enforced epigenetic silencing and variegation of proviral gene expression however often occur and further hamper a way toward cure of infection as well as effective application of retroviral vectors.

The site of proviral integration is one of the factors influencing the proviral expression stability (Jordan, Defechereux, and Verdin 2001; Skupsky et al. 2010; Senigl, Auxt, and Hejnar 2012; H.-C. Chen et al. 2017; Miklík, Šenigl, and Hejnar 2018; Vansant et al. 2020). Retroviral integration is not restricted to particular genomic positions and occurs genome-wide. Locally, the integration is affected by imperfect integrase preferences for DNA composition projected into a weak mixed motif at target sequences (Miklík et al. 2023). Globally, the integration is affected by preferences for chromatin modifications projected into skewed genera-specific genome-wide proviral integration distribution (Mitchell et al. 2004; Schröder et al. 2002; Wu et al. 2003; Narezkina et al. 2004; LaFave et al. 2014; De Ravin et al. 2014; Elleder et al. 2002; Derse et al. 2007). It is therefore often challenging to define whether a retrovirus is sensitive to the particular epigenomic environment in a given cell type or whether the observed proviral expression instability is the function of stochastic noise in proviral gene expression (Burnett et al. 2009; Weinberger et al. 2005; Miller-Jensen et al. 2013). The question of whether and how the environment at the site of proviral integration affects proviral expression becomes important in clinics when i) proviral expression is challenged to cure a latent reservoir of infection, ii) stable transgene expression is required after gene delivery in cell and gene therapies.

The effect of the integration site environment on proviral expression was demonstrated in several retroviral systems. Proviruses of avian sarcoma and leukosis virus (ASLV), that are effectively silenced in mammalian cells, were shown to be protected from the silencing when integrated closely downstream from active promoters (Senigl, Auxt, and Hejnar 2012; Šenigl et al. 2017) or close to intragenic enhancers when the vector was strengthened by insertion of the CpG island (Šenigl et al. 2017). The human immunodeficiency virus (HIV-1) expression was demonstrated to be sensitive to integration retargeting (Vranckx et al. 2016) and active HIV-1 proviruses were shown to be enriched in the vicinity of enhancers (H.-C. Chen et al. 2017; Šenigl et al. 2017; Vansant et al. 2020). Also, those silenced HIV-1 proviruses that are sensitive to latency-reversing agents tend to be integrated closer to enhancers than their reactivation-resistant (or “locked”) counterparts (H.-C. Chen et al. 2017; Battivelli et al. 2018). Active promoters and enhancers of the host genome were thus shown to act as features forming an environment permissive for retroviral transcription.

Gammaretroviruses (γRVs) target the host genome with a non-random integration distribution pattern. γRV preintegration complex interacts through the integrase C-terminal domain with the bromodomain and extraterminal domain (BET) proteins, tethering the preintegration complex to host cell chromatin (De Rijck et al. 2013; Sharma et al. 2013; Gupta et al. 2013). As a consequence, γRV proviruses are enriched at active gene promoters and enhancers (Mitchell et al. 2004; De Ravin et al. 2014; LaFave et al. 2014; Wu et al. 2003). The BET-integrase interaction can be disrupted by single point mutation (W390A) in integrase, allowing the development of BET-independent (Bin) γRV vectors that are retargeted further from promoters and enhancers (Aiyer et al. 2014; El Ashkar et al. 2014, 2017). Fusion of the chromodomain of heterochromatin-binding protein 1β (CBX) to the integrase with W390A mutation (Bin^CBX^) further reinforced the retargeting profile of the γRV vectors. In contrast to retargeted HIV-1 proviruses, the MLV Bin vector retargeting was not associated with any significant change in proviral expression and silencing. The Bin self-inactivating (SIN) vectors with heterologous internal promoters displayed stable long-term expression and minimal signs of integration site selection associated with the selection of active proviruses (Van Looveren et al. 2021).

MLV serves as a prototype virus to study the integration and expression of γRVs. Although silencing of MLV expression in murine/rat somatic cells was reported (Xu et al. 1989; Lorincz et al. 2000), we previously observed that the MLV-derived vector establishes long-term stable expression in the human K562 cell line (Miklík, Šenigl, and Hejnar 2018). Moreover, in another study, treatment of MLV-transduced HeLa cells by histone deacetylase inhibitor didn’t uncover any significant silenced proviral population (Zhu et al. 2018). Expression driven by MLV LTR was however identified as not sufficient in the human hematopoietic cells and thus, the spleen focus forming virus (SFFV) LTR was selected as a strong γRV-derived promoter for expression in hematopoietic cells (Baum et al. 1995) and has been extensively used as an internal promoter in SIN retroviral vectors (Schambach et al. 2006; Thornhill et al. 2008; Suerth et al. 2010; Warlich et al. 2011; Huston et al. 2011; Suerth et al. 2012). Indeed, SFFV LTR can drive a long-term stable expression of genes carried by MLV Bin SIN vectors (Van Looveren et al. 2021). However, the sensitivity of the SFFV LTR to epigenetic silencing in pluripotent stem cells (Hoffmann et al. 2017) together with the potent transforming activity (Modlich et al. 2009; Zychlinski et al. 2008) made the SFFV LTR unsuitable for clinical applications.

The available information suggests that γRV LTRs may act as potent drivers of expression in human somatic cells. However, the data about the expression outside the preferred loci, and the activity other than MLV and SFFV-derived LTRs is scarce. In this work, we investigated the ability of distinct γRV LTRs to establish and keep stable expression in genomic sites underrepresented in normal γRV integration. We used γRV Bin vectors and alpharetroviral (αRV) SIN vectors to study the expression and silencing of γRV LTRs after modified integration targeting. Moreover, we established CRISPR-Cas9-directed proviral knock-in method into lamina-associated domains (LADs) to demonstrate the transcriptional capacity of γRV LTRs in the transcriptionally restrictive environment.

## Materials and Methods

### Cell cultures, viral vector production and transduction

The HEK293T cell line was cultivated in DMEM:F12 medium (Sigma) supplemented with 5% fetal calf serum and 5% newborn calf serum (both GIBCO), and antibiotics (#A5955, Antibiotic Antimycotic Solution (100×), Stabilized; Sigma-Aldrich) at 37 °C and in a 5% CO_2_ atmosphere. The cell line K562 was cultivated in RPMI 1640 medium (Sigma) supplemented with 5% fetal calf serum and 5% newborn calf serum (both GIBCO) and antibiotics (#A5955, Antibiotic Antimycotic Solution (100×), Stabilized; Sigma-Aldrich) at 37 °C and in a 5% CO2 atmosphere.

Both, γRV and ASLV-derived SIN (AS).γRV vectors were produced by HEK293T cell line co-transfection. One day before the transfection, the cells were seeded on the polylysine-coated plates/dishes. The next day, cells were co-transfected by the viral genome, Gag-Pol (Schambach et al. 2006; El Ashkar et al. 2017; Suerth et al. 2012), and pVSV-G (Clontech) constructs in 6:3:1 weight ratios by X-Treme Gene HP Transfection Reagent (Roche) or the calcium phosphate method. The provider protocol was followed for transfection by X-Treme Transfection reagent. When the calcium phosphate method was used for transfection, 2.5-3 x 10^6^ cells were seeded on a p100 plate and a total of 30 μg of plasmid DNA was mixed with water (up to 1 mL) and 135 μL of 2M CaCl_2_.2H_2_O creating the mix A. The mix A was then dropwise added to the mix B formed by 1,12 mL of HBS and 22 μL of 100x concentrated PO_4_.

The complete transfection mix was added dropwise to the cells and washed after 5 h by HBS with glycerol and PBS, and the cells were supplemented by fresh cultivation medium. One day after transfection, the medium was changed for the fresh medium, and two days after transfection, the viral stocks were collected. Some viral stocks were concentrated using ultracentrifugation when the collected medium was first filtered through a 0.45 μm syringe filter (Corning) and then ultracentrifuged at 24,000 RPM using the SW40-Ti rotor for 2.5h at 4 ℃. The supernatant was then removed and the pellet was dissolved in RPMI 1640 medium (Sigma) by shaking at 4 ℃ overnight. The resulting viral stocks were stored at −80 °C.

K562 cells were counted and seeded with fresh cultivation medium on a cultivation plate/dish on the day of transduction. Virus-containing medium was added to the cultivation medium and cells were incubated for 24 hours. The next day, the medium was removed and fresh cultivation medium was added to the cells.

### Plasmids and cloning

We used pLG, the MLV-derived vector with 5’ Moloney murine sarcoma virus (MoMSV) long terminal repeat (LTR) and 3’ Moloney murine leukemia virus (MoMLV) LTR expressing enhanced green fluorescent protein (EGFP) (Kalina et al. 2007) for initial experiments with Bin vectors. For the experiments comparing LTR activity, the pLd2G vector was derived from the pLG vector by replacing EGFP with GFP fused to the destabilization domain (d2GFP). The d2GFP was amplified with Phusion Hot Start DNA Polymerase (ThermoFisher Scientific) with LdG_IF primers and inserted into EcoRI and NcoI-HF (NEB) digested pLG plasmid by In-Fusion® HD Cloning Kit (Takara) resulting in the pLd2G vector. The backbone of pLd2G was amplified by gLTR_OUT_IF primers, and the d2GFP-containing genomic sequence was amplified by gLTR_IN_IF primers. The resulting gLTR_OUT and gLTR_IN amplicons were used to create the vectors with heterologous LTRs.

MoMLV LTR was amplified from pLd2G by MoMLV_LTR primers, CrERV LTR was amplified from pCrERV-5 (Fábryová et al. 2015) by CrERV_LTR primers. Feline leukemia virus (FeLV, GenBank NC_001940 (H. Chen et al. 1998)), spleen necrosis virus (SNV, (Shimotohno, Mizutani, and Temin 1980)), and koala retrovirus (KoRV, GenBank NC_039228 (Hanger et al. 2000)) LTR sequences were synthesized as gBlocks™ Gene Fragments (Integrated DNA Technologies). LTR amplicons or gBlocks™ Gene Fragments were mixed with the pLd2G backbone (gLTR_OUT) and genomic (gLTR_IN) amplicons and joined in a single reaction using In-Fusion® HD Cloning Kit (Takara).

To construct plasmid for the Alpha.SIN.γRV.d2GFP.wPRE (AS.γRV) vector production, first, the EGFP in the original pAlpha.SIN.SF.EGFP.wPRE. plasmid (Suerth et al. 2012) was exchanged for the d2GFP by digestion with NcoI and SpeI (NEB) restriction enzymes. The d2GFP and 3’ part of the proviral genome were amplified with pAS_GFP or pAS_PRE primers. The digested backbone and the two amplicons were joined by In-Fusion® HD Cloning Kit (Takara). The SFFV U3 in the pAS.SFFV.d2GFP.PRE plasmid was then exchanged for other U3 sequences by digesting the plasmid with XcmI and AscI (NEB) restriction enzymes. The U3 parts were amplified by pAS_*γRV*U3_InFu primers, the upstream and downstream U3-flanking sequences were amplified withpAS_XcmI, pAS_5intP and pAS_AscI-intP primers. Resulting amplicons were mixed with the digested backbone and joined in a single reaction by In-Fusion® HD Cloning Kit (Takara).

In the targeted knock-in experiments pLG-derived gamma-retroviral (MoMLV, FeLV, KoRV) and ASLV-derived pAG3 (Kalina et al. 2007) vectors have GFP replaced by mCherry, amplified by Phusion Hot Start DNA Polymerase (ThermoFisher Scientific) with gamma_mCherry_InFu or AG3_mCherry_InFu primers, in NcoI and SalI (NEB) restriction sites by In-Fusion® HD Cloning Kit (Takara) following the user guide instructions. Then, the knock-in vectors, joined by In-Fusion® HD Cloning Kit (Takara), consist of EcoRV-NdeI (NEB) pAG3 fragment as a plasmid backbone, the MfeI-EcoRV (NEB) gamma-mCherry or NaeI-EcoRV (NEB) pAG3-mCherry fragment containing proviral sequence, and homologous arms, amplified by Phusion Hot Start DNA Polymerase (ThermoFisher Scientific) with homologous arms’ cloning primers listed in the Supplementary Table, flanking the proviral parts of vectors for homologous recombination knock-in.

### Flow cytometry

Cells were placed on a 96-well plate, spun down (1800 RPM, 3 min), and resuspended in PBS with Hoechst 33258 (1000x diluted). Cells were analyzed using an LSRII (BD Biosciences) flow cytometer recording a maximum of 10,000 Hoechst-negative cells.

Before sorting, cells were spun down (10 min, 200x g) and resuspended in RPMI 1640 medium (Sigma). Cells were sorted with BD Influx (BD Biosciences) or FACSAria IIu (BD Biosciences) to cultivation medium and cultivated according to standard protocol.

### Droplet digital PCR (ddPCR)

Genomic DNA was purified by phenol-chloroform extraction and diluted with water to a concentration of around 10 ng / μl. One μl of genomic DNA (10 ng) was added to a 20 μl reaction. Primers were added in the final concentration of 200 nM, probes with the concentration of 100 nM. Reactions were prepared in duplicates. Droplets were generated with 20 μl of reaction mix and 70 μl of oil with a QX200 Droplet Generator. Samples (40 μl) were transferred to 96 well plates. The amplification program for probes was run as follows: 95 °C 5 min; 40 cycles of 94 °C 30 s, 58 °C 30 s, 72 °C 30 s; 98 °C 10 min. The amplification program for EvaGreen mix was: 95 °C 5 min; 40 cycles of 95 °C 30 s, 60 °C 1 min; 4 °C 5 min; 90 °C 5 min. Both programs were run with a ramp rate of 2 °C / s. QX200 droplet reader was used to read the samples.

Proviral copy number quantification was performed with ddPCR™ Supermix for Probes (BioRad), primers targeting proviral sequence, primers targeting endogenous RPP30 sequence, and probes with FAM (provirus) or HEX (RPP30). q5GFP primers/probe were used to quantify GFP-encoding proviruses, and q3mCh primers/probe were used to quantify copy number of mCherry-encoding proviruses. Droplet Generation Oil for Probes was used to generate droplets. Copy number of provirus per cell was calculated as a copy number of FAM-positive target / 2x HEX-positive target.

Purified genomic DNA from K562 cells was used for the quantification of copy number of loci targeted by guide RNAs. QX200™ ddPCR™ EvaGreen Supermix was used in the reaction mix together with the primers used for the target site amplification / chromatin immunoprecipitation (ChIP). QX200™ Droplet Generation Oil for EvaGreen was used to generate droplets.

### gRNA design and CRISPR-Cas9 cleavage efficiency

For the design of gRNAs, we used the Crispor program (http://crispor.tefor.net/) (Concordet and Haeussler 2018). Because the CRISPR-Cas9 vector used in this study is pSpCas9(BB)-2A-GFP (PX458) (#48138, Addgene) with U6 promoter, so, we added G nucleotide at the 5’ end of the sense oligo (A) and C nucleotide at the 3’ end of the antisense oligo (B) if necessary. gRNAs oligo-duplexes were ligated into the BbsI excision site during the simultaneous digestion-ligation step. Therefore, CACC and CAAA are added to the 5’ end of the sense and the antisense oligos, respectively (Ran et al. 2013). Oligonucleotide sequences of gRNA oligos are listed in the Supplementary Table. The gRNAs named DoT:1-5.x are designed into randomly selected intra-LAD B2 and B3 subcompartments, and gRNAs named DoT:6-7.x are designed into intra-LAD B2 and B3 subcompartments from the dataset of MLV integration sites obtained in transduction experiment Figure 1D. Next, we designed gRNA, called reg9, into a previously targeted domain (Tasan et al. 2018). We also used IFT20-gRNA2 targeting the active gene IFT20 (Katoh et al. 2017).

**Figure 1.**
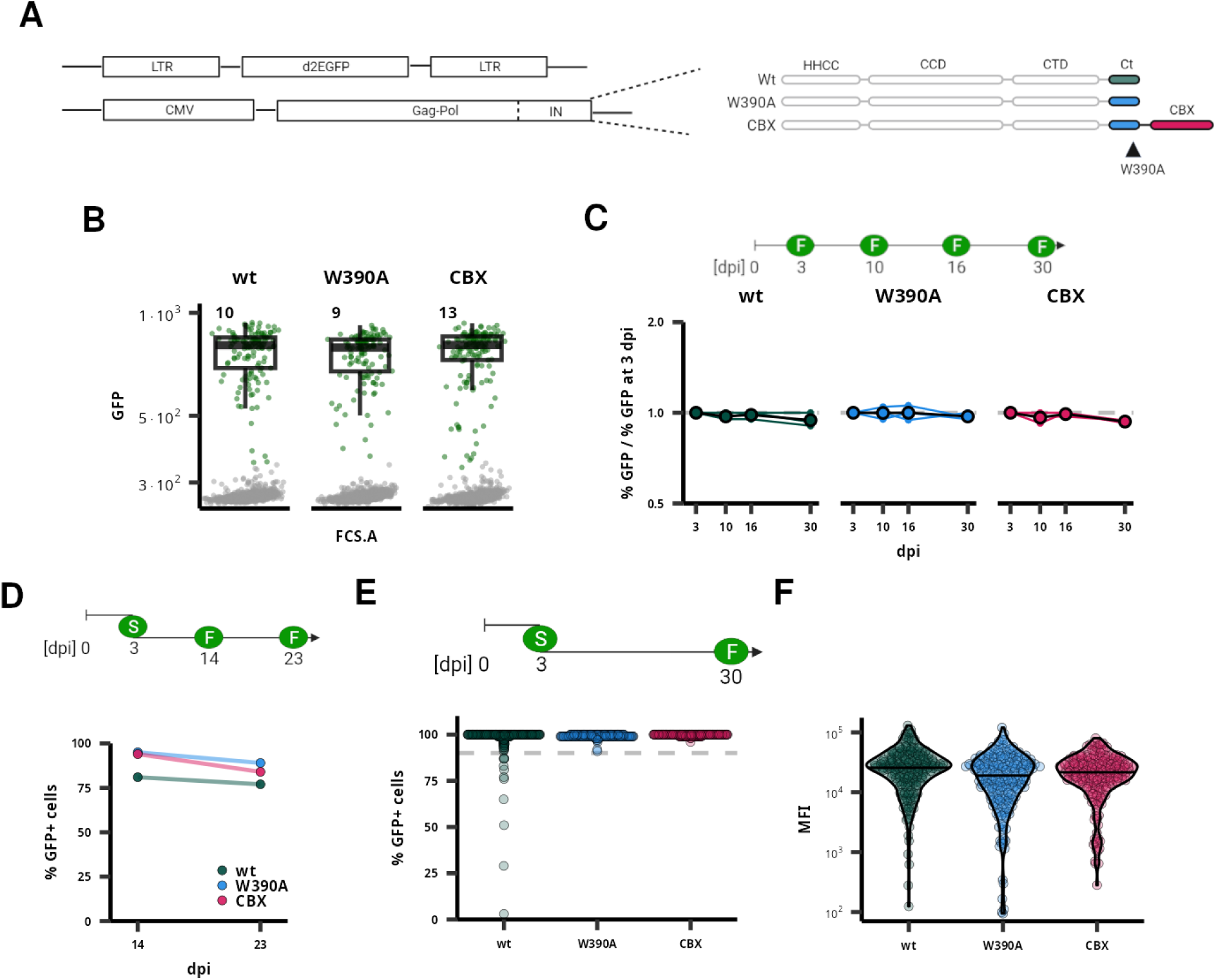
MLV is stably expressed after integration retargeting. Comparison of expression stability of MLV-derived vectors after integration retargeting. A) A schematic depiction of the LTR-dGFP-LTR vectors and integrase (IN) variants used in the experiment. B) A dot plot representing the flow cytometry measurement of K562 cells transduced by MLV-derived vectors 3 dpi. Numbers correspond to the percentage of GFP+ cells. For each vector variant, 2400 live cells were selected to construct the dot plot. C) Fold change in the fraction of GFP+ cells in the transduced population during 30 days of culture. The y-axis is on a log_2_ scale. Timepoint 3 dpi represents the data in panel B. D) Fraction of GFP+ cells after the cultivation of bulk populations sorted for GFP expression at 3 dpi. E) and F) demonstrate characteristics of clonal populations expanded from single cells sorted for GFP expression at 3 dpi. The fraction of cells expressing GFP E) and the mean fluorescence intensity (MFI) F) were measured at 30 dpi. In each category, 201 clonal populations were characterized. Schemes in panels C), D), and E) show a time course of experiments with flow cytometry (F) or FACS sorting (S) performed at a given dpi.

To test the CRISPR-Cas9 cleavage efficiency, we transfected K562 cells with 3 µg of pX458 vector with gRNA. Two days after transfection, we sorted 3000 GFP+ cells into 100 µl of lysis buffer (10 mM Tris-HCl, pH 8.0, 1 mM EDTA, 0.2 mM CaCl2, 0.001% Triton X-100, 0.001% SDS, 1 mg/ml proteinase K) per sample. After incubation of cells in the lysis buffer at 58 °C for 1 hour, and then 95 °C for 10 min, the target site was amplified by Taq DNA polymerase:DeepVent polymerase 200:1 U (both NEB) with CRISPR efficiency primers and sequenced. Then, the ratios of indels in CRISPR-Cas9 treated samples to the non-treated sample were determined in ICE CRISPR Analysis Tool (https://ice.synthego.com/<u>#/</u>; Synthego Performance Analysis, ICE Analysis. 2019. v3.0. Synthego; [2022]).

The indel profile of every gRNA site was then used for homologous arms design with homologous arms immediately adjacent to the cleavage site (Supplementary Figure S14). Individual homologous arms differ in length. We tried to design approximately 300-1000 bp long homologous arms. Nevertheless, we were limited by repetitive sequences to design unique primers. The lengths of homologous arms are stated in the Supplementary Table.

### Targeted knock-in

The K562 cells were co-nucleofected with 3 µg of each vector, pX458 with gRNA and the LTR-driven mCherry expressing vector with homologous arms to the target sites, in the Nucleofector™ I/II/2b Device (Lonza). None of the vectors were linearized prior to nucleofection. Two days post nucleofection, we sorted the cells for GFP, and 9 days post nucleofection, we sorted the mCherry+ cells in a single-cell mode. Expanded clones had mCherry expression measured by flow cytometry (LSRII) 30-46 dpn and DNA isolated on the columns (DNA Blood and Tissue Kit, Qiagen).

Obtained clones were tested for targeted insertion by amplification by BioTherm DNA polymerase (GeneCraft) with intra-proviral and homologous arms-adjacent sequence primers listed in the Supplementary Table. Only clones verified from both upstream and downstream genome-insert junctions were further analyzed for mCherry copy number by ddPCR (CN < 1.5 = single copy per genome, CN ≥ 2.5 = 3 or more copies per genome; Biorad). Then clones with 2 copies per genome had the target site amplified by BioTherm DNA polymerase (GeneCraft) with target site amplification primers listed in the Supplementary Table. Triploid DoT:7.3 should have one remaining empty allele amplified, while other target sites should be targeted and therefore not amplified. Clones were analyzed in batches with master mixes. GAPDH sequence was used as a qualitative control of amplified DNA.

### DNA library

The DNA library was prepared as described in (Hron, Fabryova, and Elleder 2020) with slight modifications. Briefly, after phenol-chloroform DNA extraction from transduced cells, DNA was fragmented by DNA fragmentase (NEB) to obtain fragments 100-2000 bp long with the main density between 500-1000 bp on an agarose gel. By purification fragments on SPRI Magnetic beads (Canvax), we selected fragments 300-2000 bp long. Then, we repaired the ends and added 3’polyA overhangs with T4 polymerase, T4 polynucleotide kinase, and Taq DNA polymerase (all NEB). After purification with the High Pure PCR Cleanup Micro Kit (Roche), through T-A ligation, we ligated adaptors with Quick Ligase (NEB). Apart from the standard adaptor oligos Ion-Link-A and Ion-Link-Ba, we added blocking oligo Ion-Link-Bb to prevent adaptor-adaptor amplification. After another purification on columns, DNA was amplified with adaptor-specific and provirus-specific primers. In the PCR reaction mix, we added blocking oligos to prevent the amplification of inner proviral sequences. First, we ran a linear PCR reaction with biotinylated primer MLV-LTR1_F_Bio. After purification of the PCR reaction on the Dynabeads™ MyOne™ Streptavidin C1 beads (Invitrogen), we ran a PCR reaction with the Ion_trP1 primer and the barcoded IonA_MIDx_mlvLTR2 primers. We isolated 300-500-bp-long PCR products from an agarose gel. Then, sequencing adaptors were added. Samples were sequenced on the Ion Torrent platform. The oligonucleotides are listed in the Supplementary Table.

### Integration sites

First, reads in the FASTQ file were sorted according to the barcodes using the *cutadapt -g ^B --overlap 8 --discard-untrimmed* command (Martin 2011) where *B* stands for the barcode sequence, and the read names were modified to contain the name of the barcode group. Reads containing a sequence of LTRs were sequentially selected first using *cutadapt -g ^GCTTGCCAAACCTACAGGTG --overlap 20 --discard-untrimmed* command to select reads containing a sequence targeted by LTR-specific primer. Next, reads containing the last 9 nucleotides of the LTR sequence with residual sequence of the minimal length of 15 bp were selected using *cutadapt -g ^GGTCTTTCA --overlap 8 --discard-untrimmed --minimum-length 15* command. Adapter sequences were then removed from the reads with *cutadapt -a ACCACTAGTGTCGAC --overlap 10 --minimum-length 15* command and the reads containing amplified inner proviral sequence were removed using the *cutadapt -g TTCCCCCCTT --overlap 10 --discard-trimmed* command.

Trimmed FASTQ reads were mapped to both hg19 and hg38 human genome assemblies using the Bowtie 2 (Langmead and Salzberg 2012) with *bowtie2 -p 20 -q -x hgX* command where the *hgX* marks the name of the assembly. Reads that map from the start of the read (*“MD:Z:0”* reads) that show single hit in the genome (“*XS:i:*” reads) were selected and converted to BED file using *samtools view -S -b* (Danecek et al. 2021) and *bedtools bamtobed -cigar -i (Quinlan and Hall 2010)* commands. Each proviral integration site (IS) is recorded as a single genomic LTR-proximal position. ISs supported by at least 5 reads were selected. If more ISs were present within the distance of 5 bp, only the IS supported by the most reads were selected. The two IS sets (IS_hg19.bed and IS_hg38.bed) were used independently against the annotations of respective genomic assemblies.

The set of random genomic ISs was created by the concatenation of three IS files and subsequent usage of *bedtools shuffle* command.

### Distance to features

We obtained the following annotations from publicly available databases: the genomic segments (Ernst and Kellis 2010) from the University of California Santa Cruz (UCSC) genome annotation database (https://hgdownload.cse.ucsc.edu/goldenpath/hg19/database/wgEncodeAwgSegmentationChro mhmmK562.txt); the chromatin subcompartments (Rao et al. 2014) from gene expression omnibus (SE63525_K562_Arrowhead_domainlist.txt.gz); the lamina-associated domain (LAD) genomic coordinates from 4D Nucleome Data Portal (Dekker et al. 2017; Reiff et al. 2022) (data accession 4DNFIV776O7C). Distance to the features was calculated using *bedtools closest -d* command.

### Software and statistical analysis

The cytometric data were gated using FlowJo™ v 10.10 Software (BD Life Sciences) and gated populations were exported to csv files. The exported data were subsequently concatenated and analyzed with custom R code (R Core Team 2022). All statistical tests and plots were produced with R. Figures were produced with the ggplot2 package (Wickham 2016). The impact effect-size analysis was performed with the ImpactEffectsize package (Lötsch and Ultsch 2020). The schemes in the figures were created with BioRender (https://www.biorender.com/).

### Crosslinking Chromatin Immunoprecipitation

ChIP was performed according to the slightly modified protocol described in (Alinikula et al. 2011). Briefly, after crosslinking, chromatin was sheared in a sonicator (Bioruptor, Diagenode) to obtain fragments with largest density between 100-1000 bp. Sonicated chromatin was immunoprecipitated (Anti-Histone H3 [ab1791], Anti-Human IgG [ab2410] and Anti-Lamin B1 [ab133741]; abcam), washed, and de-crosslinked. DNA was purified with High Pure PCR Product Purification Kit (Roche) and analyzed by Digital Droplet PCR (Biorad). Reactions were run in QX200™ ddPCR™ EvaGreen Supermix with ChIP primers in the following program: 95 °C 5 min; 40 cycles of 95 °C 30 s, 59 °C 20 s and 72 °C 30 s; 4 °C 5 min; 90 °C 5 min.

## RESULTS

### MLV promoter drives stable transgene expression of Bin vectors

In our previous work, we showed that cells transduced by MLV-derived vectors express the transgene stably for two months (Miklík, Šenigl, and Hejnar 2018). Also, Van Looveren and colleagues showed that promoters derived from spleen focus forming virus (SFFV) long terminal repeat (LTR), and the eukaryotic translation elongation factor 1 α (EF1α) establish long-term stable expression of the transgene in the context of Bin vectors. Here we adapted an identical approach to test whether the disruption of the natural promoter/enhancer preference of MLV integration also disrupts the ability of MLV LTRs to drive high, long-term stable expression. We utilized a previously used MLV-derived mini-vector LG (Kalina et al. 2007) and produced the vectors with wt integrase (IN^wt^) or Bin vectors with W390A mutation (Bin^W390A^) or with fused CBX chromodomain (Bin^CBX^) (Figure 1A).

First, we transduced the K562 cell line and measured the GFP expression at three days post infection (3 dpi) by flow cytometry. All vectors efficiently expressed GFP with no marked differences in the intensity of expression (Figure 1B, Supplementary Figure S1A). Next, we followed the long-term stability of expression in the population of transduced cells. We observed that the provirus-expressing cells form a stable part of the population for at least 30 dpi, irrespective of the vector used or the multiplicity of infection (Figure 1C, Supplementary Figure S1B). We also followed proviral expression in the polyclonal GFP+ populations bulk-sorted at 3 dpi (Figure 1D). The GFP+ cells formed a stable part of the cell population. At 23 dpi, we observed the lowest GFP+ fraction in IN^wt^ vector-transduced cells. However, the GFP+ cells still formed more than 75% of the population and this fraction has been observed unchanged since 14 dpi.

Finally, we examined the expression in the clonal populations expanded from the low multiplicity transduced single GFP+ cells sorted at 3 dpi. We characterized the expression of 200 single-cell clones in each vector-defined group at 30 dpi and observed that in all three groups, the majority of the clones contained more than 90% of GFP+ cells (Figure 1E). The intensity of GFP expression was not significantly changed in the clonal populations transduced with Bin vectors (Figure 1F). Some expression instability was observed only among the clones transduced by the IN^wt^ vector where 11 clones (5.5%) showed less than 90% of GFP+ cells from which 2 clones contained even less than 50% of GFP+ cells. No clones with less than 90% GFP+ cells were observed among those transduced with Bin vectors. The data thus suggest that the MLV proviruses that are active at 3 dpi are resistant to gene expression silencing for at least one month irrespective of the integrase variant used.

### Expressed MLV proviruses show a retargeted integration site profile

The Bin vectors were reported to integrate less effectively into traditional MLV sites marked by open chromatin histone modifications with a shift toward silent heterochromatin (El Ashkar et al. 2017). This shift in the integration site (IS) distribution of Bin vectors was observed in both active and non-selected proviral populations (Van Looveren et al. 2021). Since we observed no differences in the expression stability between MLV IN^wt^ and Bin vectors, we investigated whether the expected differences in IS distribution are preserved in the populations of expressed proviruses. We sorted the polyclonal populations of GFP+ cells at 3 dpi and analyzed the distribution of proviral ISs according to known genomic and epigenomic features.

As the Bin^CBX^ vector was reported to display the most significantly retargeted profile, we first tested whether there is any significant difference in proviral IS distribution between the active proviruses of IN^wt^ and Bin^CBX^ vectors. First, we calculated the distances to the nearest genome segment (Ernst and Kellis 2010) for each IS and compared the distance distributions between the vector groups. We used the “Impact” effect-size analysis (Lötsch and Ultsch 2020) to quantify the segment-relative distance differences between the distribution of IN^wt^ and Bin^CBX^ ISs. While there is a low Impact observed for the majority of sequences (68% with absolute value of Impact below 0.3), there are several segments with medium to high Impact (above 0.5) (Figure 2A, Supplementary Figure S2A). High Impact was associated with central tendency difference (CTDiff) as well as the difference in distribution shape (Supplementary Figure S2B) of proviral distance distribution relative to the genome segments associated with the active transcriptional start sites (Tss, TssF), enhancers (Enh, EnhF), and the segment associated with the 5’ end of transcription units (Gen5) (Figure 2A). The common characteristics of those segment-relative distance distributions is that the median distance to the features is very low and is gradually increased in both Bin vector ISs with the Bin^CBX^ vector ISs displaying the highest median distance to the segments (Figure 2B, Supplementary Figure S2C). For instance, the median distance to the Tss segment was 0.8 kb, 11.4 kb, and 22.6 kb in the IN^wt^, Bin^W390A^, and Bin^CBX^ IS data sets. The median IS distance to the Enh segment increased from 1.8 kb in the IN^wt^ to 8.5 kb and 15.6 kb in the Bin^W390A^ and Bin^CBX^ IS data sets. On the other hand, there were no or very marginal changes in distribution toward the segments associated with weak enhancers (EnhW, EnhWF) and DNase hypersensitive sites (DNaseD, DNaseU, FaireW).

**Figure 2.**
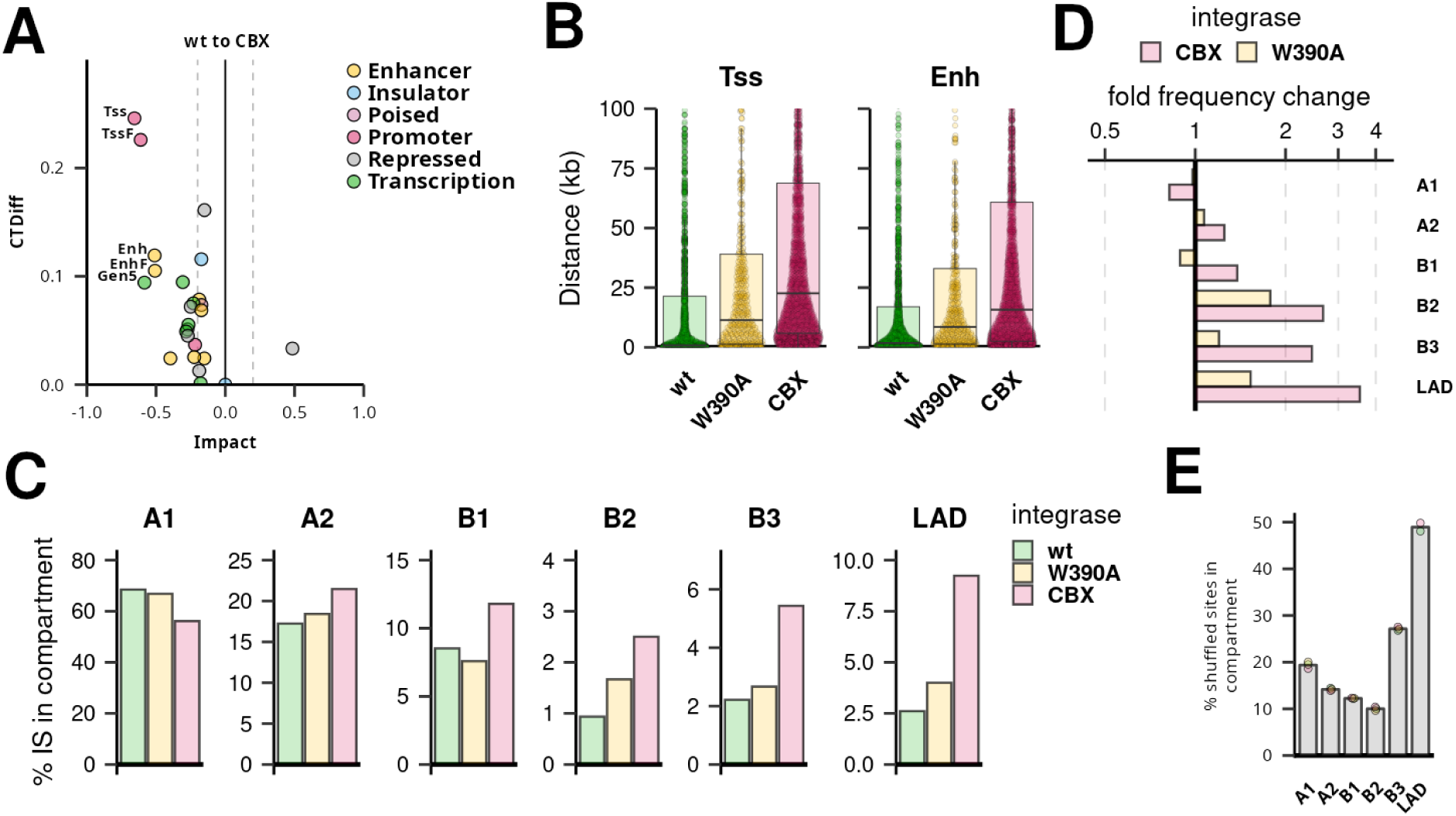
Integration site (IS) profile of active MLV proviruses after retargeting. IS distribution analysis of MLV IN^wt^ and Bin vectors in K562 cells expressing GFP. Panels A) and B) represent an analysis of IS distances to defined chromatin segments. A) Dot plot representing the results of “Impact” effect-size analysis comparing IS of Bin^CBX^ and IN^wt^ vector usage on the distribution of ISs. Points represent individual segments grouped into categories differentiated by colors. CTDiff marks the change in the central tendency of the distribution. Shown are the names of the segments with Impact absolute value ≥ 0.5. B) Plot showing the distribution of IS distances to the active transcription start site (Tss) and strong enhancer (Enh) chromatin segments. Each dot represents an individual IS, box plots represent medians and quartile range of the distance distributions. C), D), and E) panels represent the targeting of chromatin A/B subcompartments and lamina-associated domains (LAD). C) A barplot representing a fraction of proviruses integrated into the subcompartments and LADs. Fold frequency changes in subcompartment targeting of W390A and CBX IN variants to wt IN. The x-axis is depicted in the log_2_ scale. E) Frequency of shuffled sites in the chromatin subcompartments representing a random targeting control. Each dot represents a shuffled site set prepared for each of the IN variant samples. The height of the bar represents the mean targeting frequency.

Next, we tested whether the active proviruses would show altered distribution with respect to the larger domains associated with transcriptionally active or repressed chromatin. We collected the chromatin A/B subcompartments and lamina-associated domains (LADs) position data and analyzed the frequency of proviruses in those domains. The majority of proviruses were found in the A subcompartments (Figure 2C). Specifically, 68% and 17% of the IN^wt^ ISs were found in the A1 and A2 subcompartments. While the Bin^W390A^ ISs resembled the IN^wt^ ISs in A subcompartment targeting, the Bin^CBX^ ISs were found within the A1 subcompartment with decreased frequency (56%) and were increased in the A2 subcompartments (21%). Despite this minor shift, the Bin^CBX^ ISs were still significantly enriched in the A1 subcompartments compared to the random control sites (Figure 2E).

In contrast to the A compartments, the B compartments and LADs represent the infrequent locations of the active MLV proviruses. The B1, B2, B3 subcompartments and LADs contained 8%, 1%, 2.2%, and 2.6% of the IN^wt^ ISs. Similarly to the A subcompartments, the Bin^W390A^ ISs showed only a moderate change compared to the IN^wt^ IS distribution. However, the Bin^CBX^ ISs were enriched more than two-fold in the B2 and B3 subcompartments and LADs (Figure 2D). The highest enrichment was observed in the LADs where almost 9% of active Bin^CBX^ ISs were found (Figure 2C, D). Despite this increase, this frequency was still far below the expected 50% produced by random genome targeting (Figure 2E).

Here, we showed that the stably active MLV proviruses of Bin vectors display an altered profile of IS distribution. We found the active MLV proviruses to be positioned further away from the IN^wt^-preferred TSS and enhancers and more frequently situated in the heterochromatin-associated compartments.

### The stable expression after integration retargeting is common to non-MLV gammaretroviruses

The previous results suggest that the MLV-derived vectors can establish and maintain stable expression in a non-preferred epi/genomic environment. We further asked if this feature is common to other γRVs or is specific to MLV. We constructed gammaretroviral mini-vectors where the GFP with fused destabilization domain (d2GFP) transcription is controlled by the LTRs derived from Moloney murine leukemia virus (MoMLV), feline leukemia virus (FeLV), spleen necrosis virus (SNV), koala retrovirus (KoRV), and cervid endogenous retrovirus (CrERV) (Figure 3A). All constructs are capable of LTR-driven expression of d2GFP after transfection (Supplementary Figure S3). Except for the MoMLV LTR that displays a lower intensity of post-transfection expression, other LTRs drive the d2GFP expression to the intensities comparable to the expression controlled by the CMV promoter. We thus produced the vectors for the d2GFP transduction with the MLV-based packaging system utilizing the IN^wt^, Bin^W390A^, and Bin^CBX^ variants and evaluated expression stability in the transduced K562 cell line.

**Figure 3.**
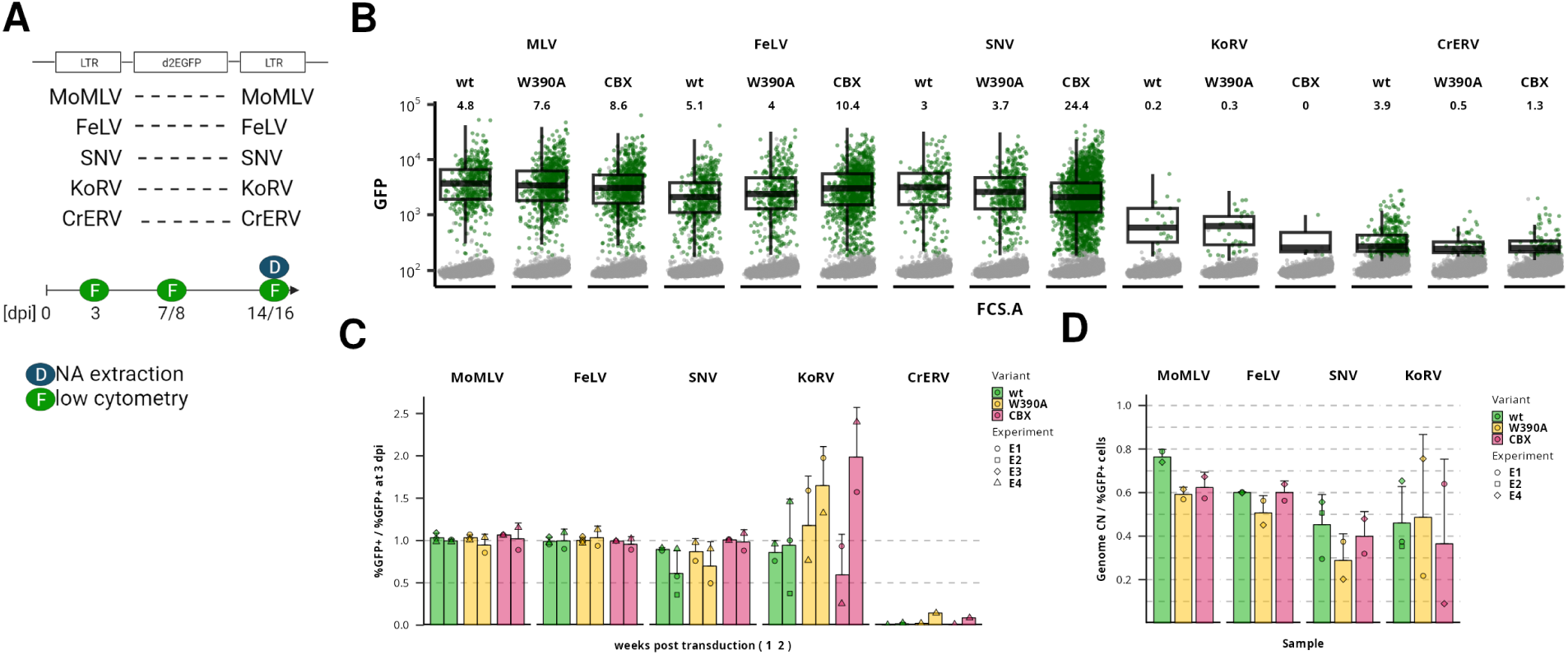
Expression stability of non-MLV gammaretroviral vectors after retargeting. A) Gammaretroviral vectors carrying LTRs from five different retroviruses were constructed. Viral stocks were produced with Gag-Pol variants and expression of the proviruses in transduced cell population was observed for two weeks. B) Expression of d2GFP by gammaretroviral vector-transduced cells at 3 dpi. For each sample, 10,000 cells are shown. GFP-positive cells are displayed in green. Boxplots show the median and quartile range of GFP intensity for the GFP-positive population. Numbers specify the percentage of GFP-positive cells in the transduced population. Shown data represent experiment E4. GFP and FSC. Log_10_ transformed GFP and FSC. A signal is used in a graph. C) Barplot showing the change in % of GFP-positive cells in time. The values are relative to the level of expression observed at 3 dpi of a particular experiment. D) Ratio of % GFP-positive cells to a copy number of detected d2GFP-encoding genomes per 100 cells. The flow cytometry and DNA extraction were performed two weeks after transduction.

At 3 dpi, we observed vector expression in the transduced cells of each vector group (Figure 3B). The FeLV- and SNV-derived vectors expressed GFP to an extent similar to the MoMLV-derived vector. On the other hand, transduction by the KoRV- and CrERV-derived vectors was less efficient; transduction by the KoRV-derived vector produced a low number of low-intensity vector-expressing cells, the CrERV-derived vector expression was dim and close to the intensity of the negative population. When cells were transduced with the IN^wt^ vectors, the FeLV-vector transduced cells displayed 1.8 and 1.5-fold lower median of GFP intensity than the MoMLV- and SNV-transduced cells, respectively. Although the transduction efficiency does not seem to be altered in the Bin vector-transduced cells, we observed slight shifts in the expression intensity in the GFP+ cells. We observed a decrease in the median GFP fluorescence intensity in the MoMLV and SNV-derived Bin^CBX^ vector-transduced cells compared to the IN^wt^ variant (1.2 and 1.5-fold, respectively, Supplementary Table S1). In contrast, using the FeLV-derived vector, we observed a 1.4-fold increase in expression intensity in cells transduced by the Bin^CBX^ variant compared to the IN^wt^ variant. In fact, the median expression intensity of the FeLV Bin^CBX^ vector is identical to the SNV IN^wt^ vector and only 1.2-fold lower than the MoMLV IN^wt^ vector-transduced cells. Although the shifts in the expression intensities are significant, they do not represent serious changes in the expression patterns of transduced genomes.

Next, we followed expression in the transduced cells for two weeks post infection (wpi) (Figure 3C, Supplementary Figure S4). The FeLV-derived vector displayed expression stability comparable to the stability observed with MoMLV. On the other hand, the SNV-derived vectors displayed a moderate decrease in the number of effectively transduced cells. The KoRV-derived vector-transduced cells displayed high variability of expression stability that can be attributed to the very low number of effectively transduced cells at 3 dpi. Unlike in other vector-transduced cell groups, the CrERV-transduced cells disappeared soon after the initial measurement at 3 dpi. Noteworthy, the Bin vectors could be traced over two weeks without any significant decrease in the percentage of effectively transduced cells compared to the IN^wt^ variant. We also observed no significant changes in the vector expression intensity over time (Supplementary Figure S5). Thus, although there might be slight differences between the vectors, the expression profiles were stable during the course of experiments.

Finally, we obtained the per-genome copy number of GFP in the transduced polyclonal populations at 2 wpi. We then related the percentage of GFP+ cells in the population to the genome copy number to estimate a fraction of expressed proviruses (Figure 3D, Supplementary Figure S6). In all samples, the ratio was below 0.8 suggesting that a population of non-expressed vectors is present in each vector-transduced population. The ratio was highest in the MoMLV-IN^wt^-transduced cells and slightly decreased to 0.6 in the MoMLV Bin^W390A^, MoMLV Bin^CBX^, and all FeLV-transduced cells. In the SNV-transduced cells, the ratio ranged between 0.3 to 0.5, which would suggest that more than half of integrated vectors with the SNV LTRs are silent at 2 wpi. Similar ratios were observed in the KoRV-derived vector transduced cells, although due to a low number of GFP+ cells, there are high deviations between data from different experiments. Generally, low copy numbers of vector genomes were observed in the CrERV-derived vector-transduced cells suggesting that the loss of actively transduced cells is possibly due to the defect in integration but not due to expression silencing.

Here, we showed that the gammaretroviral mini-vectors equipped with FeLV, SNV, and KoRV LTRs can be produced with the MLV-derived packaging system and that they can be used for stable transduction of human cells. The usage of an alternative Bin packaging system did not abrogate the stability of transgene expression in none of the vector systems. The FeLV- and SNV-derived vectors proved to be capable of expressing GFP with high intensity while the KoRV-derived vectors produced only a few low-expression transduced cells. Noteworthy, the FeLV-derived Bin^CBX^ vector showed efficient vector production and transduction efficiency comparable to the MLV-derived vectors.

### Gammaretroviral LTRs as internal promoters in αRV SIN vector

Previously, we studied the activity of the γRV LTRs in the context of γRV minigenome vectors derived from the MLV packaging system. However, we were able to accomplish effective cell transduction only with MoMLV, FeLV, and SNV-derived vectors. To further investigate the activity of all studied γRV LTRs, we utilized αRV ASLV-derived SIN (AS) vectors (Suerth et al. 2012). First, the integration of αRV vectors has no preference for any specific genomic feature (Mitchell et al. 2004; Narezkina et al. 2004), second, the transgene is expressed from the internal promoter that can be derived from retroviral LTR. We inserted the 3’ unique (U3) parts of the LTRs of MoMLV, FeLV, SNV, CrERV, or KoRV into the AS vector originally equipped with the promoter/enhancer derived from SFFV and compared the post-transduction expression stability of the derived AS.γRV.d2GFP vectors in the K562 cell line.

All AS.γRV.d2GFP vectors effectively transduced K562 cells producing up to 40% GFP+ cells at 3 dpi (Supplementary Figure S7A). Analysis of the expression intensities of the GFP+ cells revealed significant differences between the vectors (Figure 4B, Supplementary Table S3). Cells transduced with AS.MoMLV, AS.SFFV, and AS.FeLV vectors reached the highest intensities among the AS.γRV vectors. The median intensity of GFP expression was about 2-fold lower in cells transduced with AS.SNV and AS.CrERV vectors (Supplementary Table S3). The AS.KoRV-transduced cells were present with the lowest expression intensities, 5-fold lower than the AS.MoMLV and 2-fold lower than the AS.SNV vector-transduced cells.

**Figure 4.**
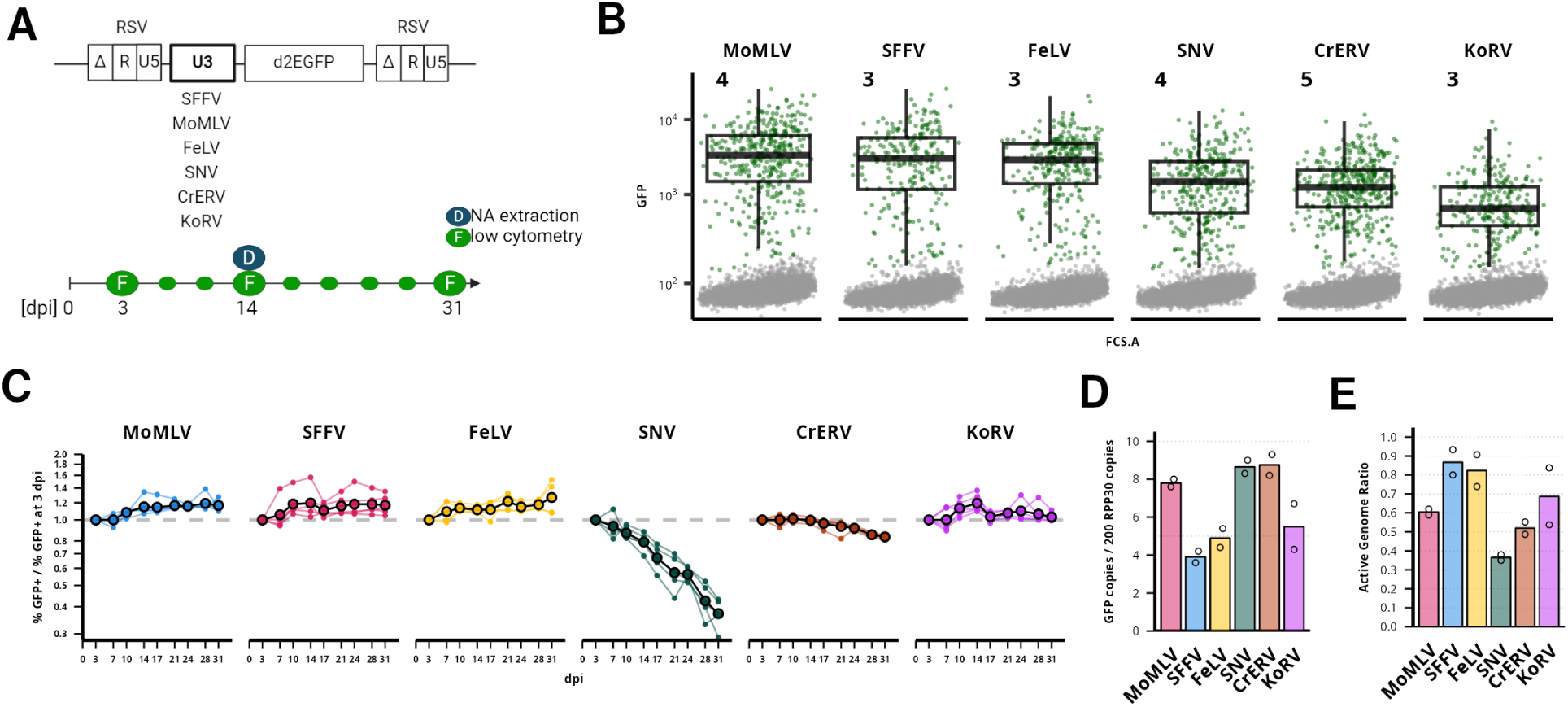
Intensity and stability of αRV vector expression with gammaretroviral LTR as an internal promoter. A) A schematic depiction of an alpharetroviral (αRV) AS vector. The internal promoter was derived from the U3 promoter/enhancer part of LTR of studied gammaretroviruses. The bottom of the panel contains the scheme of the experiment where GFP expression was followed every 3 - 4 days 3 - 31 dpi. B) Intensity of d2GFP expression in transduced K562 cells at 3 dpi. Cells inside the GFP+ gate are colored green. Boxplot describes the intensity distribution of GFP+ cells. The numbers show the percentage of GFP+ cells of all alive cells. For each sample, 10,000 cells are shown. Samples are ordered by the median intensity of GFP+ cells. C) Representation of the time-course experiment where the percentage of GFP+ cells in transduced populations was followed. Values on the y-axis show the percentage of GFP+ cells relative to the percentage of GFP+ cells observed at 3 dpi. Light lines and points show individual transduction experiments with divergent multiplicities of infection. Black-outlined points connected by black lines show the average of all experiments. D) Copy number of proviruses (GFP copies) as per 100 genomic equivalents (200 copies of RPP30 reference target) measured by the droplet digital PCR (ddPCR). Proviral copy number was established from genomic DNA collected at 14 dpi in samples shown in panel B. E) Ratio of the percentage of GFP+ cells per proviral copy number per 100 genome equivalents. The value of 1 marks the point where all proviruses are expected to be active in expression. Points in D) and E) show values of technical duplicates.

Next, we examined the proportion of GFP+ cells in the transduced populations every 3 to 4 days for up to 31 dpi (Figure 4C). The proportion of GFP+ cells in AS.MoMLV, AS.SFFV, and AS.FeLV-transduced populations first slightly increased after 3 dpi and stayed stable since. Also, AS.KoRV-transduced cells showed stable levels of GFP+ cell proportion. The proportion of GFP+ cells in AS.CrERV-transduced population was stable for about two weeks after transduction, but then started to slowly decrease at 17 dpi and at 31 dpi reached about 83.5% (±1.6%) of the proportions observed at 3 dpi. We observed a rapid decrease in the proportion of GFP+ cells in the AS.SNV-transduced cells (Supplementary Figure S7B). At 31 dpi, the mean GFP+ cell proportion reached 37.5% (±5.6%) of the proportions observed at 3 dpi.

We collected genomic DNA from the low-multiplicity transduced populations at 14 dpi and performed ddPCR to estimate the proviral copy number in each vector-transduced population. At 14 dpi, the proportion of the GFP+ cells was comparable in all selected populations (mean 3.9% ±0.6%). The levels of proviral copy number, however, fluctuated between 3.9 (AS.SFFV) to 8.8 (AS.CrERV) proviral copies per 100 equivalents of cellular genomes (Figure 4D). We then estimated the active genomes ratio and observed that AS.SFFV together with AS.FeLV are the most active vectors producing 80 to 90% of active proviruses at 14 dpi. AS.MoMLV and AS.KoRV displayed lower proportion of active genomes between 60 to 70%, the ratio observed with Bin vectors. We observed the lowest proportion of active genomes (36%) in AS.SNV-transduced cells. As the overall proportion of GFP+ cells decreases over time, we estimate that already at 3 dpi, the proportion of active genomes of AS.SNV is 46% and may go down to 13% at 31 dpi (Supplementary Figure S7C).

Here, we showed that γRV LTRs can be used as internal promoters in SIN vectors. In the context of αRV SIN vector, MoMLV and FeLV LTRs produce high-intensity and stable expression of the vector at levels comparable to the SFFV LTR. However, contrary to MoMLV LTR, FeLV LTR produces low frequency of proviruses silenced early after integration. KoRV LTR also produced a long-term stable proportion of vector-expressing cells albeit with much lower expression intensity than other vectors.

### Gammaretroviruses are stably expressed after targeted knock-in into LADs

In infection and retargeting experiments, we revealed the LADs integrations in the population of active γRV proviruses. To test the expression capacity of γRV proviruses in the LADs directly, we utilized the CRISPR-Cas9-mediated homology-directed repair-driven knock-in method to insert retroviral mini-genomes into K562 annotated LADs. We selected the LADs containing B2 and B3 subcompartments and designed gRNAs targeting the sites covered by the subcompartments. We selected five domains randomly (domains 1-5) and two domains where the expressed MoMLV proviruses were previously localized (domains 6 and 7). We also targeted the LAD with reported vector insertion (reg9) (Tasan et al. 2018) and used previously published gRNA into the transcriptionally active gene IFT20 (Katoh et al. 2017). The coordinates of the sites targeted by gRNAs are listed in Supplementary Table S4.

First, we tested the CRISPR-Cas9 cutting efficiency in the LAD target sites. Out of the total 23 Domain-Target (DoT) sites, 4 DoTs were cleaved with relatively high efficiency: DoT:3.1 (14%), DoT:6.2 (81%), DoT:6.3 (72-74%), and DoT:7.3 (34%) (Supplementary Table S4). The characteristics of effectively cut target sites are schematically depicted in Figure 5A (see Supplementary Figure S8 for more detail). To verify the characteristics of the target sites, we examined the ploidy and the presence of Lamin B at the target sites (Supplementary Figure S9 and S10). Apart from DoT:7.3, whichshows higher copy number, other target sites are diploid. We also observed the increased levels of Lamin B at the selected DoTs relative to the active genes IFT20 and RPP30.

**Figure 5:**
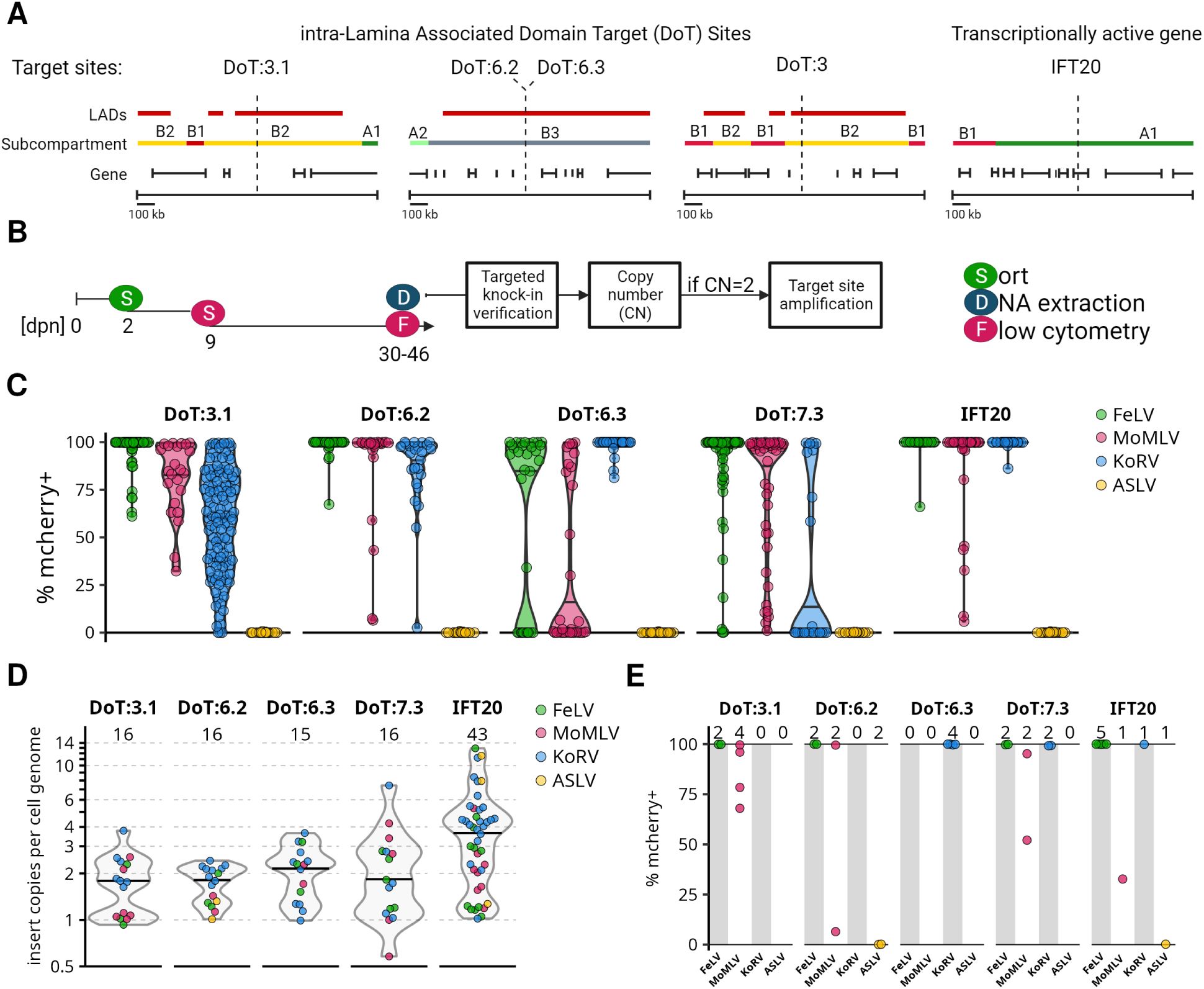
Gammaretroviral expression after insertion into Lamina-Associated Domain target (DoT) sites. A) Schematic depiction of target sites with the position of predicted LADs, chromatin subcompartments, and genes. B) The scheme of the experiment and subsequent analysis. Two dpn we sorted (marked S) GFP+ subpopulations and 9 dpn we sorted mCherry+ cells in a single-cell mode. Expanded clones were measured for mCherry expression by flow cytometry (marked F) 30-46 dpn. We also isolated DNA (marked D) from expanded uncharacterized clones and verified targeted insertions in the individual clones. Then we checked a vector copy number. If the clones contained 2 copies per genome, we amplified the target site to minimize the risk of the second copy off-target. C) Long-term proviral expression stability of uncharacterized clones in target sites 30-46 dpn. D) Determination of vector copies per genome in clones with verified targeted insertions. E) Proviral mCherry expression in clones with a verified single insertion per genome.

Next, we cotransfected the GFP-expressing CRISPR-Cas9 vectors containing selected gRNAs with the mCherry-expressing mini-vectors containing the LTRs from MoMLV, FeLV, both highly expressed after transduction, KoRV, which has low expression after transduction, and ASLV as a negative control. The promoter activity of the ASLV LTR is highly sensitive to the epigenomic environment and should be active only near active promoters (Senigl, Auxt, and Hejnar 2012; Šenigl et al. 2017). We sorted the GFP+ populations two days post nucleofection (2 dpn), and 9 dpn, we sorted the mCherry+ cells in a single-cell mode. The expanded clones were measured by flow cytometry and further characterized (Figure 5B, Supplementary Table S5).

We characterized the expression of mCherry in 788 clones at 30-46 dpn (Figure 5C). The proportion of stably expressing clones (≥ 90% mCherry+ cells) ranged broadly from 14% (DoT:3.1-KoRV) to 98% (DoT:6.2-FeLV) among the populations with the DoTs insertion. Among the clones where DoT:3.1, DoT:6.2, and DoT:7.3 sites were targeted, the vector with the FeLV LTRs displayed the most stable expression producing 90, 98, and 73% of stably expressing clones. In contrast, among the clones where the DoT:6.3 site was targeted, the vectors with the KoRV LTRs displayed the most stable expression with 95% stably expressing clones. The clones containing the MoMLV-derived vector displayed intermediate stability of expression in each of the DoT group. We observed the highest stability of the MoMLV-derived vector among the clones in the DoT:6.2 site group producing 83% of stably expressing clones. We saw less striking differences in vector expression among the clonal populations where the IFT20 gene was targeted. In this case, 80, 97, and 97% of clones with MoMLV, KoRV, and FeLV-derived vectors, respectively, were stably expressing the mCherry after more than one month of cultivation. Noteworthy, expression observed in the clones where a MoMLV-derived vector was targeted to IFT20 resembles the expression stability observed after MoMLV transduction (Figure 1E). In comparison to γRVs, the introduction of ASLV-derived vectors produced no clones displaying more than 1% of mCherry+ cells.

We next tested 301 clones for a full-length homology recombination-mediated provirus knock-in by amplification of the upstream and downstream genome-insert junctions. We verified the targeted insertion in 106 clones. The expression profile of provirus-expressing clones did not differ from the previous population of expanded clones (Supplementary Figure S11). We were able to verify insertion into target sites only in 5 ASLV-derived clones. However, we verified the targeted insertion of ASLV-derived vectors in mCherry-negative populations (Supplementary Figure S12). Next, we determined the mCherry copy number in the verified clones by ddPCR (Figure 5D). We observed a single copy in 30 clones, two copies in 35 clones, and in 41 clones we observed 3 or more copies of the mCherry. A median mCherry copy number per genome in DoT insertion-verified clones was around 2, while the IFT20-targeted clones’ median copies per genome was 3.5 with some clones reaching more than 10 mCherry copies per genome.

We obtained clones with a verified single insertion in every DoT site, but only DoT:7.3 was represented by all 3 γRV vectors (Figure 5E). Irrespective of a target site, most proviruses were stably expressed for at least 30 dpn. We also analyzed the clones with 2 copies per genome, which have a higher risk of off-targets, but likely contain each copy of the vector inserted in a separated allele of the target site (Supplementary Figure S13). In two-copy clones, only the DoT:3.1 was represented by all 3 γRV vectors. The target site IFT20 was not verified in this group of clones. From 10 double-insertion clones, most clones were stably expressing mCherry, however, MoMLV-a KoRV-containing clones were unstable in the DoT:3.1.

Here, we reported the targeted insertion of retroviral mini-genomes into B2/B3 subcompartments of LADs. We demonstrated that γRV vectors can be long-term expressed in LADs and that the environment of individual DoT sites functionally differ in the strength of the support of γRV expression.

## Discussion

The genomic and epigenomic environment at the site of proviral integration is one of the key determinants influencing the expression of the provirus. Since γRVs present a sharp integration preference into active promoters and enhancers, the features associated with the protection of retroviral transcription (Senigl, Auxt, and Hejnar 2012; Šenigl et al. 2017; Vansant et al. 2020; H.-C. Chen et al. 2017), we examined whether promoter activity of γRV LTR is dependent on the environment preferred by γRV integration. Our results demonstrate that despite a significant proportion of proviruses being silenced early after integration, distinct γRV LTRs establish and maintain long-term stable expression after the disruption of the natural integration preference. Moreover, we directly demonstrate the ability of γRVs to establish stable expression in a restrictive environment by direct knock-in of γRV proviruses into LADs.

First, we quantified the postintegration expression stability of γRV LTRs. We observed only limited expression silencing or variegation of γRV LTR-driven gene expression after 3 dpi not only with Bin vectors but also with γRV LTRs as internal promoters in the αRV SIN vector. This observation is consistent with previous reports using the IN^wt^ MLV-derived vector (Miklík, Šenigl, and Hejnar 2018) or Bin vectors with heterologous internal promoters (Van Looveren et al. 2021). We observed strong silencing after 3 dpi only with SNV LTR as internal promoter in the αRV SIN vector. The RU5 part of the SNV LTR that is missing in the internal SNV promoter was shown to act as a positive postranscriptional control element enhancing cytoplasmic expression of RNA (Butsch et al. 1999; T. M. Roberts and Boris-Lawrie 2000; Tiffiney M. Roberts and Boris-Lawrie 2003). Our results suggest that the RU5 region may also play a positive role in transcription stability. Although rarely observed, the variegation of expression is most frequent in the MLV vector with natural integration preference. Notably, the variegation of MLV expression is observed only when expression of individual proviruses is tracked in clonal populations. Presence of variegating MLV proviruses thus seems to depend on integration site selection, although, unexpectedly, the variegation is observed in the population of proviruses with the natural preference for promoters and enhancers.

Next, we observed that with normal integration targeting in the K562 cell line, about 20% of MLV proviruses are silent early after infection, i.e. before the markers of proviral expression are first detected (3 dpi in our study). The silent population can consist of both epigenetically silenced and damaged proviruses (Xu et al. 1989; Lorincz et al. 2000). Our observation that the proportion of silent proviruses doubles after integration retargeting with Bin vectors supports the existence of epigenetically silenced population and further supports that the concept where promoter activity of MLV LTR is to some extent sensitive to the epigenetic environment at target site. Missing expression variegation and the increase in silent population of MLV proviruses after retargeting suggests that BET-directed integration targets MLV to specific, semi-permissive sites missed by Bin vectors. The phenomenon of increased proportion of silenced population after integration retargeting seems to be specific to MLV LTR as we observed uniform proportion of silent population among wt and Bin vectors bearing FeLV and SNV LTRs. One explanation can be that, unlike MLV, FeLV and SNV LTRs are rapidly silenced at putative semi-permissive sites. Our results show that significant transcriptional silencing of all tested γRV LTRs occurs rapidly after proviral integration. The proviruses that escape the early silencing events then perform with the long-term stable expression. The early post-integration silencing of γRV LTRs in somatic cells is a phenomenon that should be addressed in future studies.

We showed that the natural preference of γRV directed by BET-integrase interaction is dispensable for the establishment of long-term stable proviral expression. On the other hand, despite the retargeted proviruses being more distant from promoters and enhancers, the majority of retargeted active proviruses we characterized are still positioned inside predicted chromatin A compartments and can thus be supported by the transcriptionally permissive environment of the chromatin domain. The observed retargeted IS profile can, however, still in part result from selection pressure. However, we observed significant differences between active proviruses of IN^wt^, Bin^W390A^, and Bin^CBX^ vectors and there were only marginal differences between non-selected and active proviral populations of Bin vectors bearing internal SFFV promoter (Van Looveren et al. 2021). While these observations don’t rule out IS-specific silencing, the effect of non-permissive sites on tested γRVs LTRs is probably too weak to be detected by current bulk analysis. Some stochastic shutdown of proviral expression immediately after integration can also occur.

In vector-transduction experiments, we observed expressed MLV proviruses in LADs - a compartment generally considered to be formed by heterochromatin. However, some endogenous genes in LADs can be transcribed (Leemans et al. 2019) and can thus form locally opened chromatin for retroviral integration and proviral expression. We thus sought to insert proviruses into selected intergenic, B2/B3 subcompartment-associated intra-LAD regions by CRISPR-Cas9-mediated homology-directed repair. We successfully inserted MoMLV, FeLV, and KoRV LTR-containing proviruses into LADs and observed that the LTRs can drive transgene expression inside LADs. Most importantly, we demonstrated that stable long-term transgene expression can be achieved with a single provirus inserted at the different LAD target sites.

Knock-in by CRISPR-Cas9-mediated homology-directed repair is commonly applied to insert transgenes into selected genomic loci like *AAVS1*, *CCR5,* or *TRAC* (Eyquem et al. 2017; Lombardo et al. 2007, 2011). Such target sites are situated inside transcribed genes which allows for insertion of promoterless cassettes expressed under the control of endogenous promoters. Effective expression in LADs, however, would require the utilization of silencing-resistant promoters. We demonstrated that γRV LTRs can serve as such promoters. However, the expression heterogeneity between different LTRs in different LAD target site that aligns well with the notion of heterogenic environment of intra-LAD regions (Leemans et al. 2019) suggests that proper site-promoter combination should be tailored for effective cassette expression in the LAD target sites. Targeting the DoTs resulted in fewer inserted proviral copies per cellular genome than targeting the actively transcribed gene IFT20. One possible explanation may be the formation of cassette concatemers at the target site (Lombardo et al., 2007) that may be more frequent in transcribed loci. Therefore, targeting the LAD target sites may lead to a lower copy number of inserted expression cassettes, which could result in better control over the outcomes of gene knock-in.

While designing gRNAs for DoTs we avoided possibly active regions by omitting intragenic regions and selecting B2/B3 subcompartments. We then selected only those sites showing indels introduced by error-prone pathway repairing CRISPR-induced double strand breaks. As only few of the selected DoTs showed indels, the sites probably differ in the accessibility of DNA for cleavage or by kinetics of the repair mechanisms (Brinkman et al. 2018). Although recombination by homology directed repair was reported inefficient in LADs (Tasan et al. 2018), any DoTs showing even small efficiency was accessible for the cassette insertion. Next, selecting LADs targeted by retroviral integration possibly increased a chance of successive CRISPR-Cas9 cleavage, even though we didn’t select the exact positions targeted by the integration. More careful post hoc inspection of successfully targeted DoTs revealed presence of additional features like predicted lncRNA or CTCF binding sites. However, features associated with targetable sites remain to be defined. We failed to detect CRISPR-Cas9 cleavage at one DoT (reg9) used for targeted cassette insertion in the HCT116 cell line (Tasan et al. 2018). Although the site is predicted to lie inside LAD in both HCT116 and K562 cell lines, the accessibility in each cell line may differ. Thus, although particular LADs may be conserved among different cell types, the accessibility of LAD target sites can be cell-type-specific.

In our study, we moved from retargeting of γRV integration to specifically targeted knock-in of proviruses to show that diverse γRV LTRs can function as promoters not only outside active promoters and enhancers but also in the silent-prone environment of LADs. This work opens the possibility of studying other than MLV γRVs as drivers of cassette expression. The presentation of the directed intra-LAD knock-in technique opens new possibilities for research of LADs as safe landing pads for targeted gene insertions.

## Supporting information

supplementary figures and data-containing tables

sequences of the oligonucleotides used in the study.

## Acknowledgement

We would like to thank Axel Schambach and Melanie Galla for sharing the wt MLV Gag-Pol construct and the constructs for production of alpharetroviral vectors, and Rik Gijsbers for sharing the modified MLV Gag-Pol constructs. We would like to thank our colleagues Lubomíra Pecnová and Markéta Reinišová for kind help during cell cultivation and PCR confirmation in the knock-in experiments.

## Data and code accessibility

Raw sequencing data, integration site coordinates and code will be accessible through public repositories during the final step of manuscript preparation.

## Supplementary data

Supplementary_Data.pdf file contains supplementary figures and data-containing tables. Supplementary_Table.xlsx file contains sequences of the oligonucleotides used in the study.

## Contributions

DM, MS, and JH designed the study parts and wrote the manuscript. JH received the funding for the project. DM performed transduction experiments, bioinformatic analysis and produced graphical representation of the data. MS prepared sequencing libraries and performed chromatin immunoprecipitation and knock-in experiments. DK cloned viral plasmids. SM and JM managed and performed sequencing of the libraries.

## Funding

This work was supported by the Czech Academy of Sciences (Praemium 677 Academiae Award 2018 and RVO: 68378050-KAV-NPUI). D.M., M.S., and J.H. were also supported by the project National Institute of Virology and Bacteriology 675 (Programme EXCELES, ID Project No. LX22NPO5103) - Funded by the European Union - Next 676 Generation EU.

## Notes

### Competing Interest Statement

The authors have declared no competing interest.

