## supplementary figures and data-containing tables for "Long Terminal Repeats of Gammaretroviruses Retain Stable Expression After Integration Retargeting or Knock-In into the Restrictive Chromatin of Lamina-Associated Domains"

SUPPLEMENTARY TABLES

SUPPLEMENTARY FIGURES

#### SUPPLEMENTARY TABLES

| | MoMLV | | | FeLV | | | SNV | | | KoRV | | | CrERV | | |
|---|---|---|---|---|---|---|---|---|---|---|---|---|---|---|---|
| | wt | W390A | CBX | wt | W390A | CBX | wt | W390A | CBX | wt | W390A | CBX | wt | W390A | CBX |
| **MoMLV_wt** | 1 | 1.1 | 1.2 | 1.8 | 1.6 | 1.2 | 1.2 | 1.4 | 1.8 | 5.7 | 5.6 | 9.4 | 9.1 | 9.9 | 9.6 |
| **MoMLV_W390A** | 1.1 | 1 | 1.1 | 1.6 | 1.4 | 1.1 | 1.1 | 1.3 | 1.6 | 5.3 | 5.1 | 8.7 | 8.4 | 9.1 | 8.8 |
| **MoMLV_CBX** | 1.2 | 1.1 | 1 | 1.5 | 1.3 | 1 | 1 | 1.2 | 1.5 | 4.7 | 4.6 | 7.8 | 7.5 | 8.2 | 7.9 |
| **FeLV_wt** | 1.8 | 1.6 | 1.5 | 1 | 1.1 | 1.4 | 1.5 | 1.2 | 1 | 3.2 | 3.1 | 5.3 | 5.1 | 5.6 | 5.4 |
| **FeLV_W390A** | 1.6 | 1.4 | 1.3 | 1.1 | 1 | 1.3 | 1.3 | 1.1 | 1.1 | 3.7 | 3.6 | 6.1 | 5.8 | 6.3 | 6.2 |
| **FeLV_CBX** | 1.2 | 1.1 | 1 | 1.4 | 1.3 | 1 | 1 | 1.2 | 1.4 | 4.7 | 4.5 | 7.7 | 7.4 | 8 | 7.8 |
| **SNV_wt** | 1.2 | 1.1 | 1 | 1.5 | 1.3 | 1 | 1 | 1.2 | 1.5 | 4.9 | 4.7 | 8 | 7.7 | 8.4 | 8.1 |
| **SNV_W390A** | 1.4 | 1.3 | 1.2 | 1.2 | 1.1 | 1.2 | 1.2 | 1 | 1.2 | 4 | 3.9 | 6.6 | 6.4 | 6.9 | 6.7 |
| **SNV_CBX** | 1.8 | 1.6 | 1.5 | 1 | 1.1 | 1.4 | 1.5 | 1.2 | 1 | 3.2 | 3.1 | 5.3 | 5.1 | 5.6 | 5.4 |
| **KoRV_wt** | 5.7 | 5.3 | 4.7 | 3.2 | 3.7 | 4.7 | 4.9 | 4 | 3.2 | 1 | 1 | 1.6 | 1.6 | 1.7 | 1.7 |
| **KoRV_W390A** | 5.6 | 5.1 | 4.6 | 3.1 | 3.6 | 4.5 | 4.7 | 3.9 | 3.1 | 1 | 1 | 1.7 | 1.6 | 1.8 | 1.7 |
| **KoRV_CBX** | 9.4 | 8.7 | 7.8 | 5.3 | 6.1 | 7.7 | 8 | 6.6 | 5.3 | 1.6 | 1.7 | 1 | 1 | 1 | 1 |
| **CrERV_wt** | 9.1 | 8.4 | 7.5 | 5.1 | 5.8 | 7.4 | 7.7 | 6.4 | 5.1 | 1.6 | 1.6 | 1 | 1 | 1.1 | 1.1 |
| **CrERV_W390A** | 9.9 | 9.1 | 8.2 | 5.6 | 6.3 | 8 | 8.4 | 6.9 | 5.6 | 1.7 | 1.8 | 1 | 1.1 | 1 | 1 |
| **CrERV_CBX** | 9.6 | 8.8 | 7.9 | 5.4 | 6.2 | 7.8 | 8.1 | 6.7 | 5.4 | 1.7 | 1.7 | 1 | 1.1 | 1 | 1 |

Supplementary Table S1. Pairwise comparison of median fluorescence intensity of transduced cells. The fluorescence intensity of K562 cells measured at 3 dpi. Vectors expressed destabilized GFP.

| -log10 P-value | MoMLV | | FeLV | | | SNV | | | KoRV | | | CrERV | | |
|---|---|---|---|---|---|---|---|---|---|---|---|---|---|---|
| | W390A | CBX | wt | W390A | CBX | wt | W390 A | CBX | wt | W390A | CBX | wt | W390A | CBX |
| MoMLV_wt | 1.8 | 14.6 | 83.8 | 36.3 | 17.4 | 7.1 | 33.9 | 135.9 | 46.8 | 78.2 | 10.6 | Inf | 141.1 | 298.7 |
| MoMLV_W390A | NA | 9.2 | 82.5 | 31.5 | 11.9 | 3.7 | 28.5 | 156.5 | 45.9 | 78.2 | 10.6 | Inf | 146 | Inf |
| MoMLV_CBX | NA | NA | 48.4 | 12.9 | 0.4 | 0.3 | 10.5 | 96.8 | 41 | 71.9 | 10.1 | Inf | 141.2 | Inf |
| FeLV_wt | NA | NA | NA | 6.9 | 44 | 29.8 | 8.4 | 0 | 28 | 54.8 | 9.4 | Inf | 128.5 | 274.2 |
| FeLV_W390A | NA | NA | NA | NA | 10.5 | 9 | 0.2 | 10.4 | 32.6 | 60.8 | 9.7 | Inf | 129.4 | 268.9 |
| FeLV_CBX | NA | NA | NA | NA | NA | 0.7 | 8.8 | 93.8 | 38.7 | 68.1 | 9.8 | Inf | 139.2 | Inf |
| SNV_wt | NA | NA | NA | NA | NA | NA | 8 | 42.6 | 36.3 | 61.9 | 9.4 | Inf | 121 | 241.3 |
| SNV_W390A | NA | NA | NA | NA | NA | NA | NA | 12.6 | 32.5 | 59.3 | 9.5 | Inf | 126.4 | 260 |
| SNV_CBX | NA | NA | NA | NA | NA | NA | NA | NA | 29.1 | 58.1 | 9.4 | Inf | 138.6 | Inf |
| KoRV_wt | NA | NA | NA | NA | NA | NA | NA | NA | NA | 0.9 | 2.5 | 17.8 | 19.3 | 22.8 |
| KoRV_W390A | NA | NA | NA | NA | NA | NA | NA | NA | NA | NA | 2.3 | 18.7 | 20.6 | 25 |
| KoRV_CBX | NA | NA | NA | NA | NA | NA | NA | NA | NA | NA | NA | 0 | 0.5 | 0.3 |
| CrERV_wt | NA | NA | NA | NA | NA | NA | NA | NA | NA | NA | NA | NA | 4.2 | 4.6 |
| CrERV_W390A | NA | NA | NA | NA | NA | NA | NA | NA | NA | NA | NA | NA | NA | 0.6 |

Supplementary Table S2. Significance of differences in pairwise fluorescence intensity comparison. The fluorescence intensity of K562 cells was measured at 3 dpi. Vectors expressed destabilized GFP. First, the Kruskall-Wallace test was run. Then the significance of pairwise comparison differences was evaluated by Wilcoxon rank sum test with continuity correction. The statistical test was run in the R environment.

**A**

| Fold Change | MoMLV | SFFV | FeLV | SNV | CrERV | KoRV |
|---|---|---|---|---|---|---|
| **MoMLV** | 1 | 1.1 | 1.2 | 2.2 | 2.7 | 5.1 |
| **SFFV** | 1.1 | 1 | 1 | 2 | 2.4 | 4.6 |
| **FeLV** | 1.2 | 1 | 1 | 1.9 | 2.3 | 4.5 |
| **SNV** | 2.2 | 2 | 1.9 | 1 | 1.2 | 2.3 |
| **CrERV** | 2.7 | 2.4 | 2.3 | 1.2 | 1 | 1.9 |
| **KoRV** | 5.1 | 4.6 | 4.5 | 2.3 | 1.9 | 1 |

**B**

| -log10 P-value | CrERV | FeLV | KoRV | MoMLV | SFFV |
|---|---|---|---|---|---|
| **FeLV** | 21.1 | NA | NA | NA | NA |
| **KoRV** | 13.4 | 35 | NA | NA | NA |
| **MoMLV** | 32 | 2.4 | 45 | NA | NA |
| **SFFV** | 16.8 | 0.4 | 28.7 | 1 | NA |
| **SNV** | 1.2 | 12.5 | 13.9 | 21.1 | 10.9 |

Supplementary Tables S3. Analysis of post-transduction expression intensities of AS.γRV.d2GFP vectors. A) Fold change of median of expression intensity of GFP+ cells. B) Statistical significance of differences in expression intensities between vectors. First, the Kruskal-Wallis rank sum test was run giving a value 77.9 (-$\log_{10}$ P-value). Then, the pairwise comparisons using Wilcoxon rank sum test with continuity correction was run. The values represent the -$\log_{10}$ P-values.

| gRNA | seq | ch | strand | start | end | cutting efficiency in K562 cells |
|---|---|---|---|---|---|---|
| DoT:1.1 | GTTGACCCACTCTTTGCACTGGG | chr10 | + | 110259331 | 110259353 | 0% indels |
| DoT:1.2 | GAGATACCACCGAGTAAATGAGG | chr10 | + | 110260355 | 110260377 | quality issues |
| DoT:1.3 | GAATAGGAGCATTGCTACATTGG | chr10 | - | 110261403 | 110261425 | quality issues |
| DoT:2.1 | GCTACTGATAGACCACATCGAGG | chr9 | + | 30194191 | 30194213 | amplification issues |
| DoT:2.2 | GATCCCCAAGACAACGGAGGAGG | chr9 | + | 30193709 | 30193731 | amplification issues |
| DoT:3.1 | GTAATCCTCTAGGGATGCCGTGG | chr20 | + | 16903611 | 16903633 | 14% indels |
| DoT:3.2 | GTCACTCAGCTATGCATAACTGG | chr20 | - | 16902842 | 16902864 | 0% indels |
| DoT:3.3 | GTGCTCCCAGGTAGCCTAGTGGG | chr20 | + | 16905773 | 16905795 | 0% indels |
| DoT:4.1 | GTGTCCTTGATATGAATCAGAGG | chr6 | - | 48479760 | 48479782 | 0% indels |
| DoT:4.2 | GAGTAGTTGACCGTAGTCATAGG | chr6 | + | 48481317 | 48481339 | amplification issues |
| DoT:4.3 | GAGTACGACTATACATATCAAGG | chr6 | - | 48481022 | 48481044 | amplification issues |
| DoT:5.1 | GCGTTGCTGCTCAGCGACTCTGG | chr20 | - | 38533862 | 38533884 | 0% indels |
| DoT:5.2 | GACAGGAAGGTATAGGCCTCTGG | chr20 | + | 38533750 | 38533772 | 0% indels |
| DoT:5.3 | GTTCCCATTGTCTAGAATCGGGG | chr20 | - | 38536522 | 38536544 | quality issues |
| DoT:5.4 | GGCAGAATGTAAGGACCGTGGGG | chr20 | + | 38538134 | 38538156 | quality issues |
| DoT:5.5 | GTTCACAAGTGTATAAGGACAGG | chr20 | - | 38538047 | 38538069 | 0% indels |
| DoT:6.1 | GATATTAGAGTATCCCGTGAG | chr12 | + | 128269430 | 128269449 | quality issues |
| DoT:6.2 | CGGTACTAAGGAGATCCCTC | chr12 | + | 128269266 | 128269285 | 81% indels |
| DoT:6.3 | ACGGTAGGAAAGCGACATTC | chr12 | + | 128268033 | 128268052 | 72-74% indels |
| DoT:7.1 | GCTAAATCCCTTGACAATTGG | chr18 | - | 1574940 | 1574959 | quality issues |
| DoT:7.2 | GTCACTCAGCACCCTATGGAC | chr18 | + | 1575126 | 1575145 | quality issues |
| DoT:7.3 | GTGTTAGCCAGCAAGTAGAA | chr18 | + | 1575252 | 1575271 | 34% indels |
| Reg9 | GCCTATACCTTTACCGATAGC | chr1 | - | 199549157 | 199549176 | 0% indels |
| IFT20 | GTGATGAACTGAACAAGCTGA | chr17 | - | 26658954 | 26658973 | N/A |

Supplementary Table S4: Coordinates of tested gRNAs (hg19). Due to repetitive elements and amplification from lysates, we did not manage to amplify some samples (amplification issues). Moreover, the sequence quality of some samples was too low to be analyzed (quality issues).

| Target site | LTR | Expanded clones | Transcriptionally stable (≥ 90 % mCh+) | Tested for knock-in | Verified knock-in | CN = 1 | CN = 2 | CN ≥ 3 |
|---|---|---|---|---|---|---|---|---|
| DoT:3.1 | MoMLV | 23 | 9 | 6 | 6 | 4 | 1 | 1 |
| | KoRV | 119 | 17 | 7 | 7 | 0 | 5 | 2 |
| | FeLV | 90 | 81 | 3 | 3 | 2 | 1 | 0 |
| | ASLV | 24 | 0 | 16 | 0 | 0 | 0 | 0 |
| DoT:6.2 | MoMLV | 23 | 19 | 3 | 2 | 2 | 0 | 0 |
| | KoRV | 24 | 14 | 15 | 9 | 0 | 8 | 0 |
| | FeLV | 40 | 39 | 4 | 3 | 2 | 1 | 0 |
| | ASLV | 24 | 0 | 16 | 2 | 2 | 0 | 0 |
| DoT:6.3 | MoMLV | 31 | 5 | 29 | 2 | 0 | 2 | 0 |
| | KoRV | 44 | 42 | 44 | 10 | 4 | 3 | 3 |
| | FeLV | 32 | 18 | 30 | 3 | 0 | 2 | 1 |
| | ASLV | 45 | 0 | 16 | 0 | 0 | 0 | 0 |
| DoT:7.3 | MoMLV | 46 | 30 | 9 | 5 | 1 | 0 | 3 |
| | KoRV | 25 | 6 | 14 | 6 | 2 | 2 | 1 |
| | FeLV | 60 | 44 | 9 | 5 | 2 | 1 | 1 |
| | ASLV | 24 | 0 | 16 | 0 | 0 | 0 | 0 |
| IFT20 | MoMLV | 30 | 24 | 11 | 8 | 1 | 5 | 2 |
| | KoRV | 30 | 29 | 24 | 21 | 1 | 2 | 16 |
| | FeLV | 30 | 29 | 13 | 11 | 4 | 0 | 5 |
| | ASLV | 24 | 0 | 16 | 3 | 1 | 0 | 1 |
| **TOTAL** | | **788** | **406** | **301** | **106** | **28** | **33** | **36** |

Supplementary Table S5: Targeted knock-in clone numbers.

### SUPPLEMENTARY FIGURES

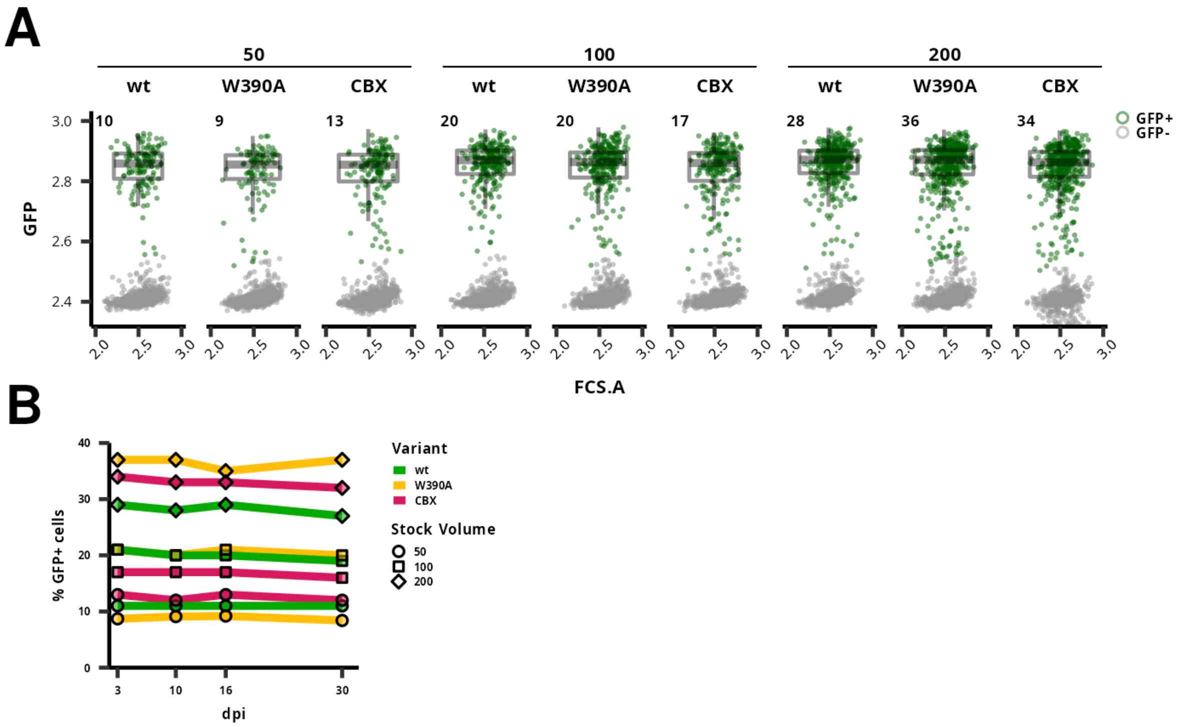


Supplementary Figure S1. A) Intensity of GFP expression of MLV-derived vector with integrase variants at 3 dpi. K562 cells were transduced by a variable volume of LG vector viral stock and GFP expression was measured by flow cytometry at 3 dpi. The numbers above the graph represent the volume of the stock added. Smaller numbers mark the percentage of GFP+ cells (in green) in the transduced populations. Box plots show the distribution of GFP intensities among the GFP+ cells. B) MLV-derived vector expression stability in time. The K562 cell line was transduced with 3 different volumes of the vector stock (in μL) represented by a point shape. Colors represent the IN variant of the vector used.

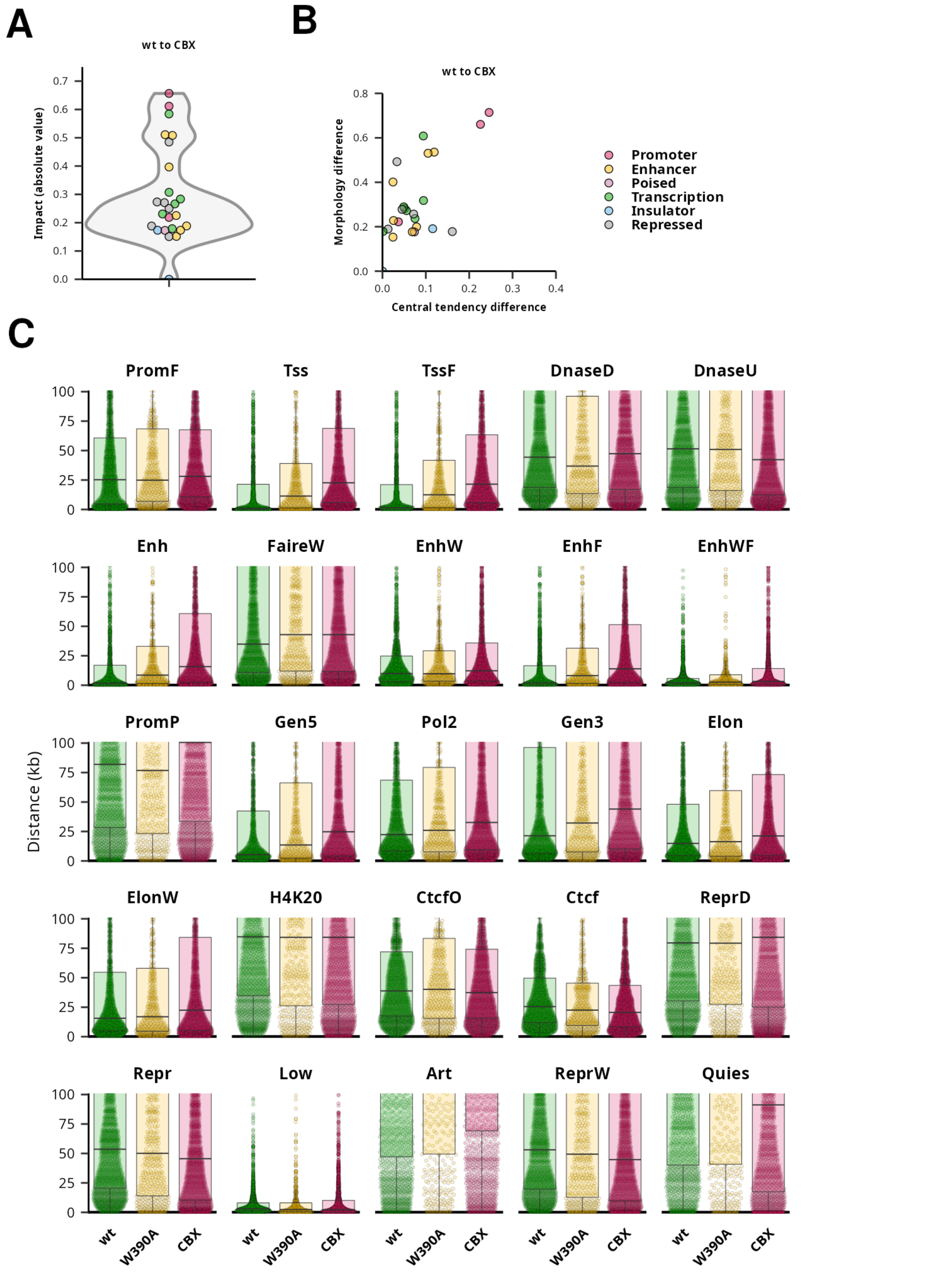


Supplementary Figure S2. Distance of proviral IS to the nearest genome segment. A) and B) Effect size analysis between IS of $IN^{wt}$ and $Bin^{CBX}$. Absolute values of Impact. B) Differences in central tendency and differences in morphology in distance distribution. C) Distances to the nearest genomic segment. Each IS is shown as a point.

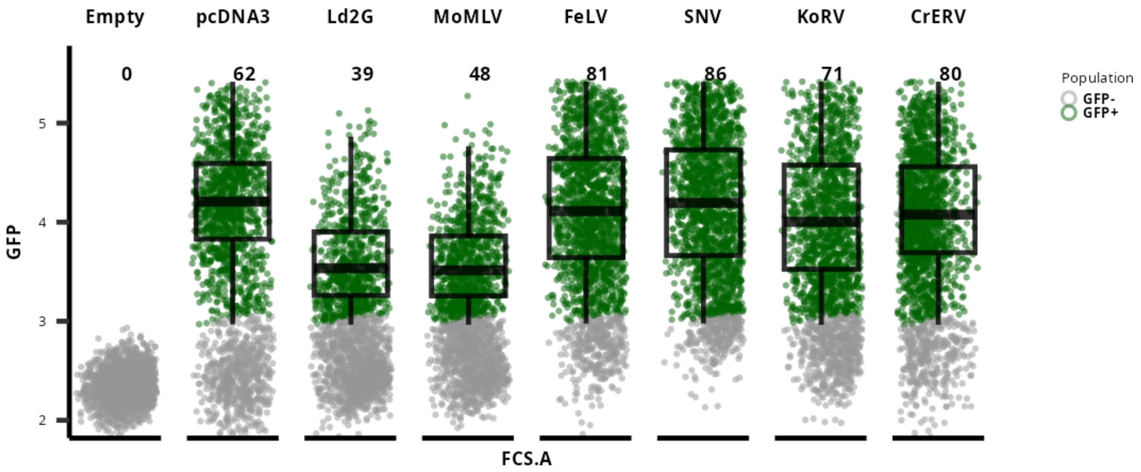


Supplementary Figure S3. Expression of destabilized GFP (d2GFP) from gammaretroviral LTRs two days after transfection of HEK293T cells. Cells were transfected on a 24-well plate with 500 µg of the plasmid DNA using X-Treme Gene HP Transfection Reagent (Roche). 2,000 Hoechst-negative cells are shown for each sample. Boxplots mark the GFP intensity distribution of cells in the GFP-positive gate. The empty vector and pcDNA3 expressing d2GFP are used as positive and negative controls. pcDNA3 uses CMV as a promoter, Ld2G is a derivative from the LG vector. Names mark the origin of the LTR in LTR-d2GFP-LTR mini-vector used for transfection and subsequent vector production. The numbers in the upper part of the plot mark the percentage of GFP-positive cells from all Hoechst-negative cells.

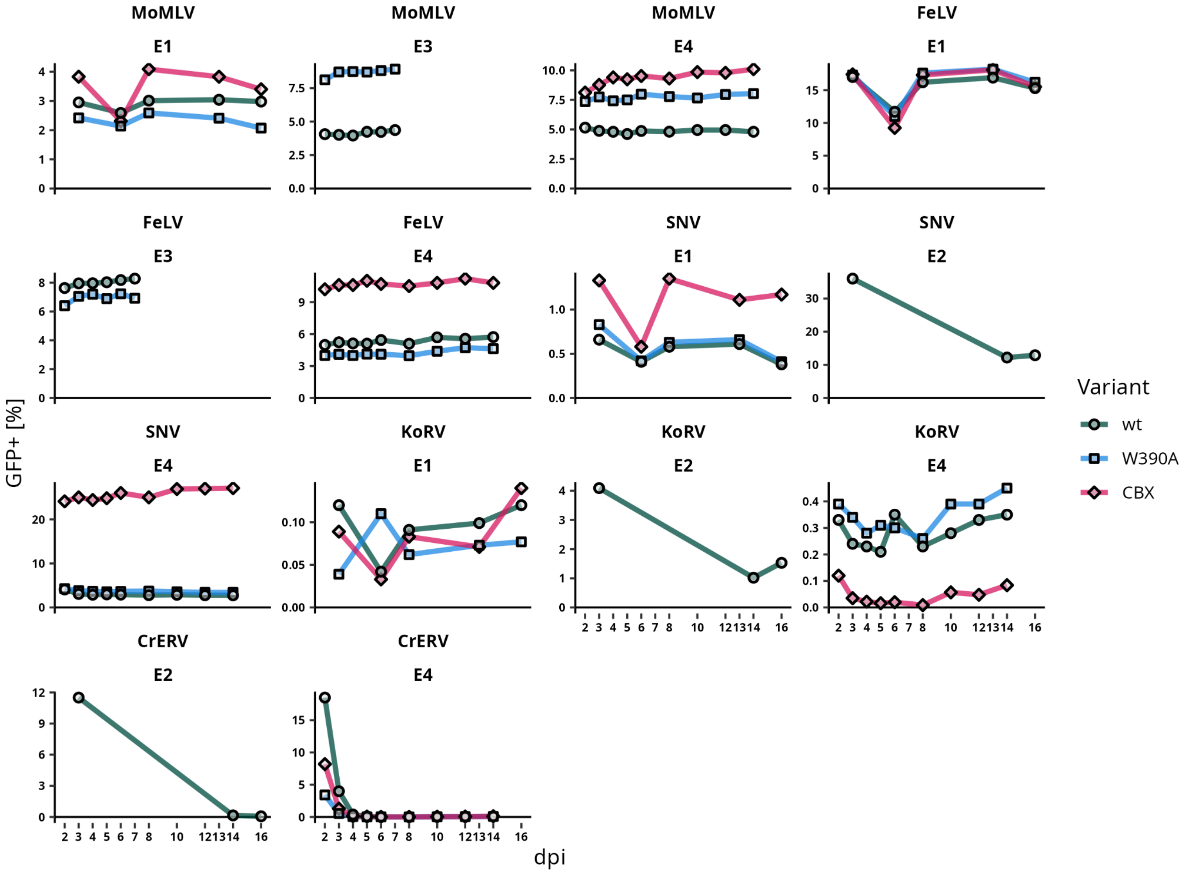


Supplementary Figure S4. Gammaretroviral vector GFP expression stability in time. Each facet represents an individual transduction experiment performed with the gammaretroviral vector. Integrase variants are distinguished by the point shape and line color.

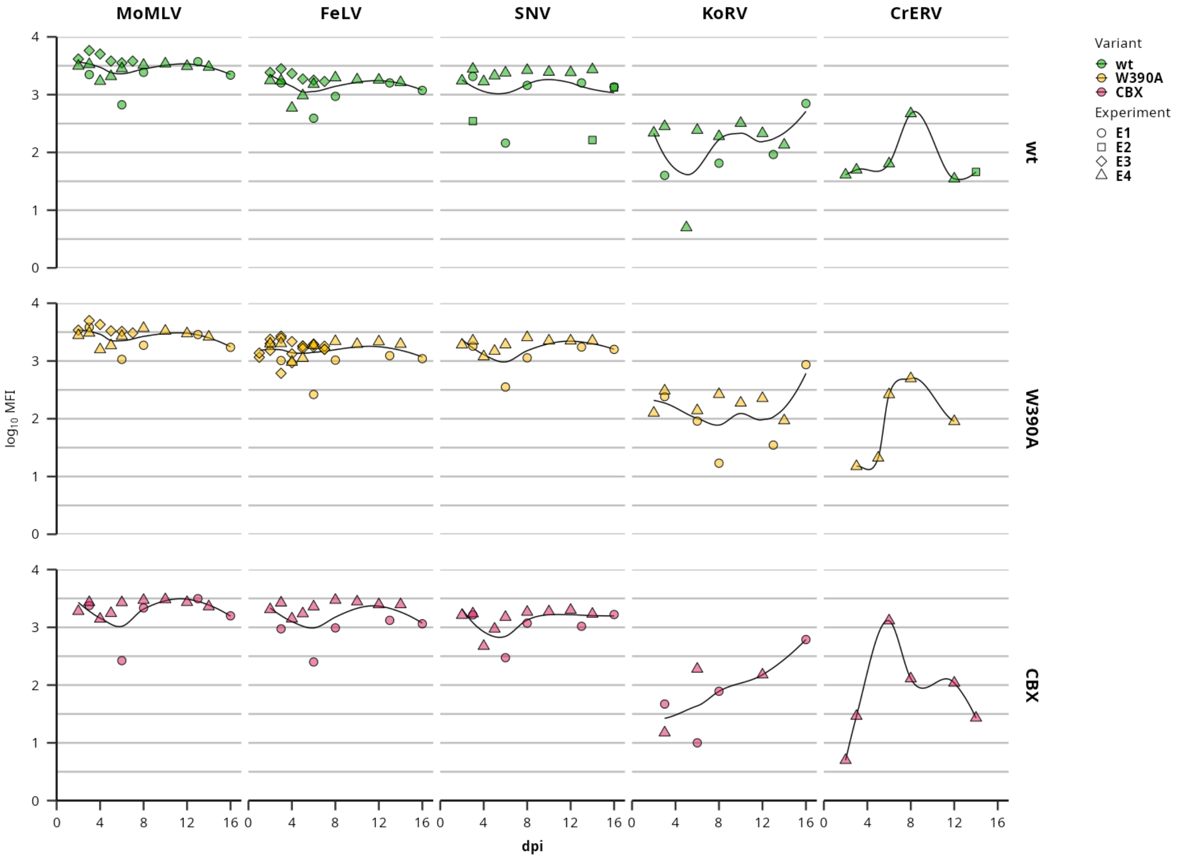


Supplementary Figure S5. Gammaretroviral vector GFP expression intensity in time. Each column represents a gammaretroviral vector, and each row represents an integrase variant. Integrase variants are distinguished by the point color, the point shape indicates the transduction experiment. The smoothed line shows the intensity trend in time. Values of the GFP-positive population mean fluorescence intensities (MFI) are shown on the $log_{10}$ scale.

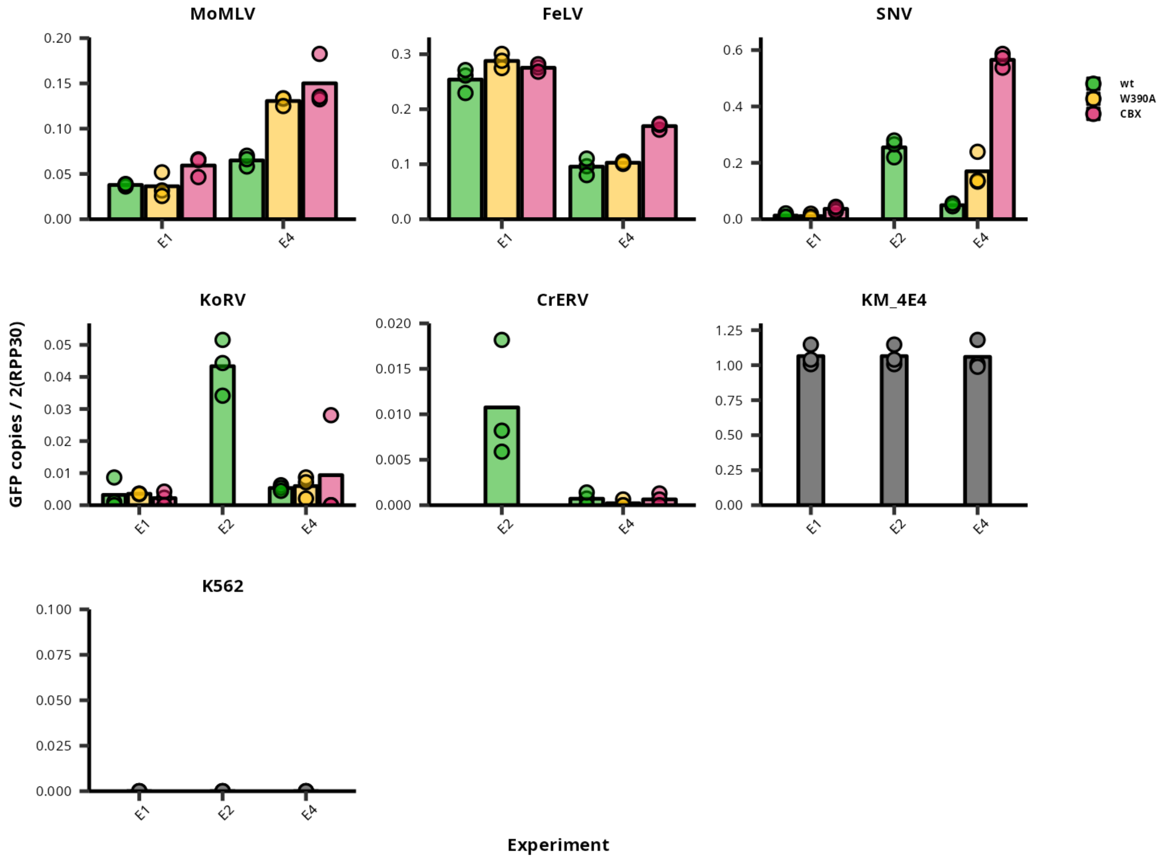


Supplementary Figure S6. Copy number of gammaretroviral vector genomes. The copy number is a value obtained as a number of GFP copies divided by a doubled number of RPP30 copies - thus receiving a mean count of GFP copies per cell genomes. Each measurement was done in technical triplicate. KM_4E4 is a genomic DNA control obtained from a cellular clone transduced by MLV-derived GFP-expressing vector containing 1 copy of vector genome per cellular genome. K562 is a control cell population of the cell line not transduced by any vector.

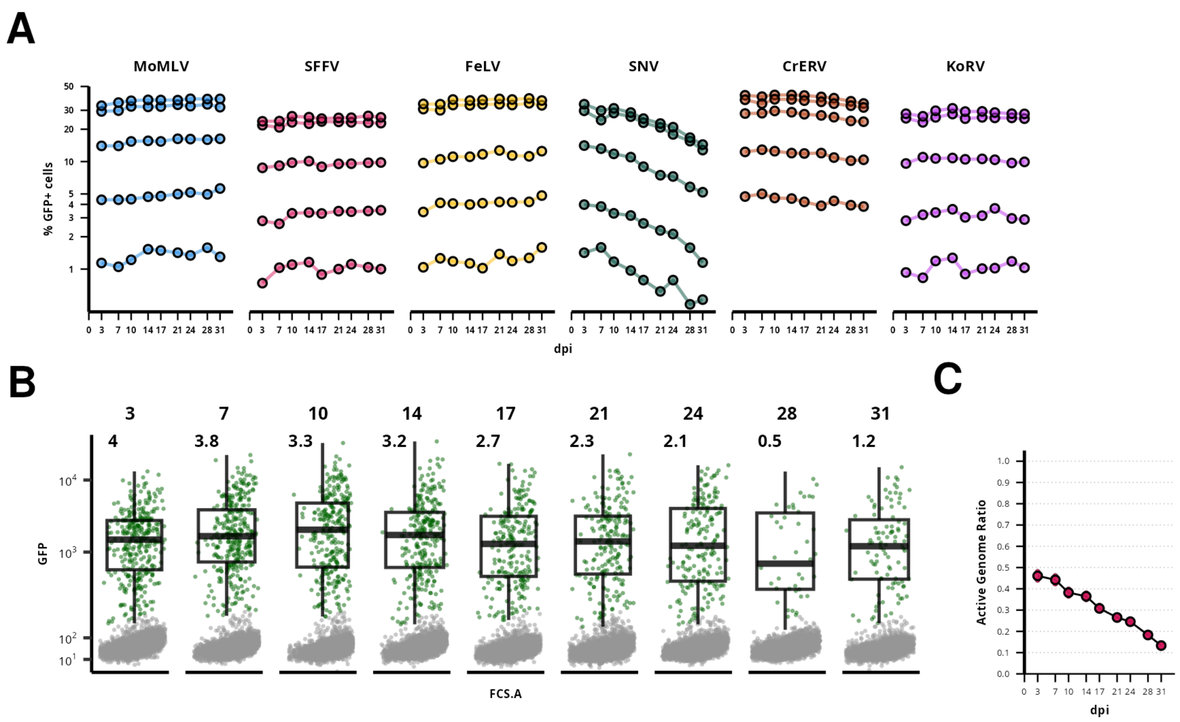


Supplementary Figure S7. Gammaretroviral LTR activity as internal promoters in the alpharetroviral vector. A) Percentage of GFP-positive cells in the K562 cell line.Each point represents a value obtained at a single measurement at a given dpi. Each line represents a single transduction experiment with various multiplicities of infection. GFP expression was followed from 3 dpi to 21 dpi.  B) Expression of the GFP by the vector with SNV LTR as an internal promoter. Upper numbers represent a dpi. Lower numbers represent a percentage of GFP-expressing cells. Boxplots represent the distribution of GFP intensity in a GFP-positive gate. C) A ratio of GFP-positive cells and GFP copies per cellular genome. The GFP copy number was quantified on genomic DNA from cells collected at 14 dpi. The active genomes ratio was then obtained as a percentage of GFP-positive cells at a given dpi divided by a number of proviral copies per hundred cellular genomes at 14 dpi.

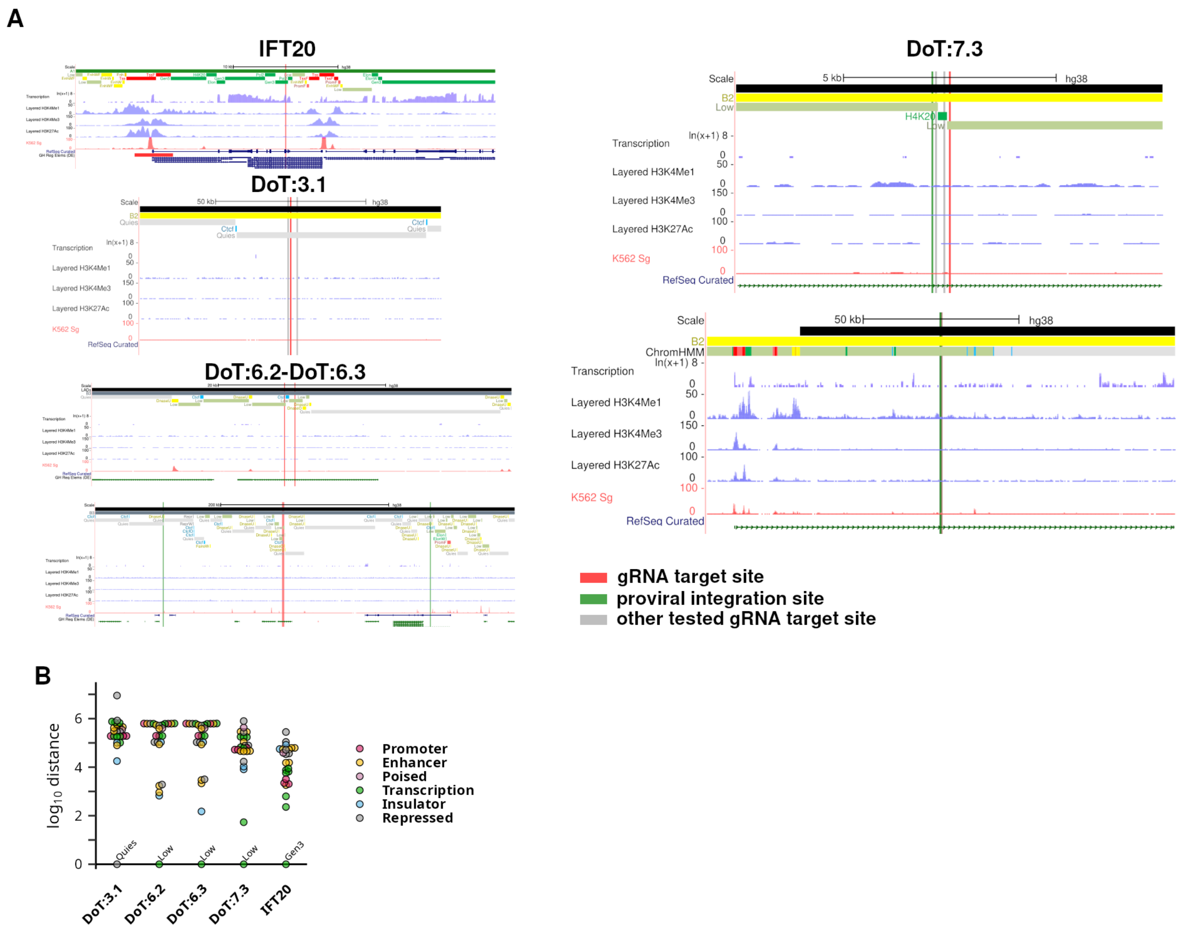


Supplementary Figure S8. The characteristics of target sites. A) The UCSC Genome Browser snapshots of the regions targeted by mini-genome insertion. Sites targeted by gRNAs are marked by red and gray lines. Sites, where proviral integration obtained from the transduction experiments was observed, are marked by green lines. B) Distance to the nearest genomic segment representing a family of genomic segments for each site targeted by mini-genome insertion.

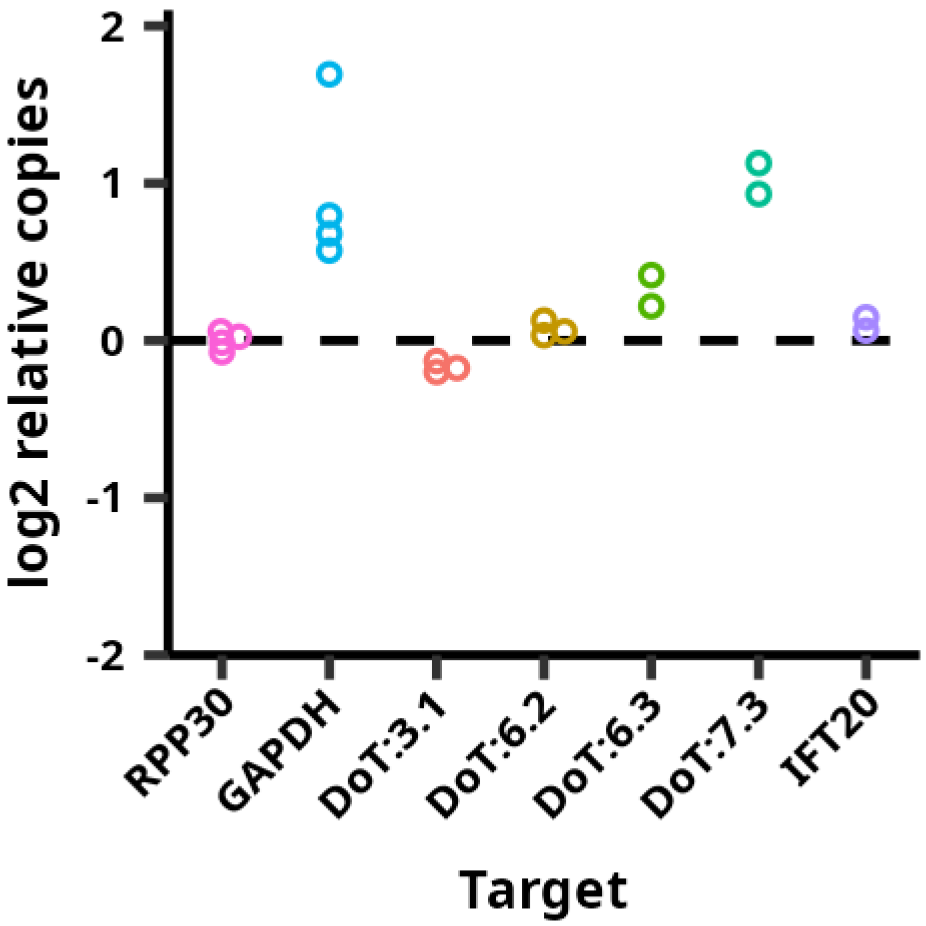


Supplementary Figure S9: Target site copy number assessed by droplet digital PCR (ddPCR) with EvaGreen. Values represent the target copy number divided by the average copy number observed for the RPP30 target region. Each dot represents a single technical replicate (a well) in the ddPCR experiment.

**A**

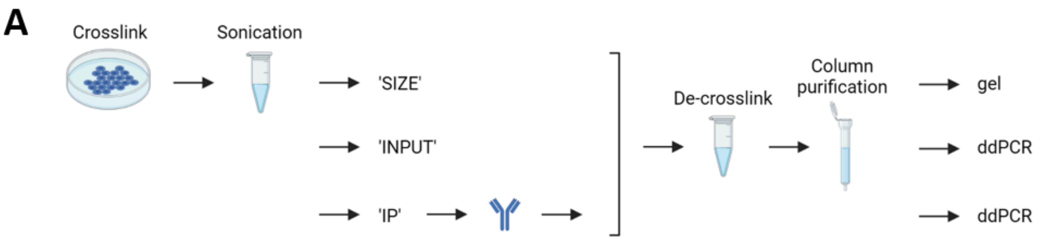


**B**

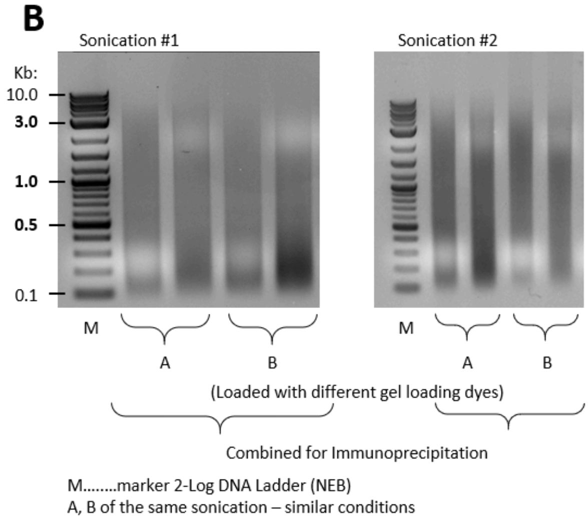


**C**

| | Anti-LaminB | | Input | | Anti-H3 | |
|---|---|---|---|---|---|---|
| DoT:3.1(1) | 3,2 | 3,5 | 52,4 | 51,5 | 119 | 119 |
| DoT:6.2(1) | 1,5 | 1,7 | 32,9 | 32,9 | 75 | 74,5 |
| DoT:6.3(1) | 1,5 | 1,7 | 33,4 | 40,9 | 78,7 | 79,3 |
| DoT:7.3(1) | 7,8 | 7,4 | 114 | 113 | 240 | 278 |
| IFT20(1) | 2,8 | 2,5 | 77,8 | 75,4 | 166 | 158 |
| RPP30(1) | 1 | 1,5 | 74,7 | 82,2 | 37,2 | 38,7 |
| DoT:3.1(2) | 12,3 | 14 | 65,2 | 76,5 | 132 | 127 |
| DoT:6.2(2) | 5,4 | 6,1 | 55,1 | 63,6 | 87 | - |
| DoT:6.3(2) | 7,3 | 7,2 | 63,5 | 64,5 | 111 | 105 |
| DoT:7.3(2) | 28,7 | 31,1 | 170 | 185 | 287 | 290 |
| IFT20(2) | 6,8 | 6,2 | 122 | 125 | 197 | 195 |
| RPP30(2) | 2,4 | 2 | 99 | 92 | 61,6 | - |

**D**

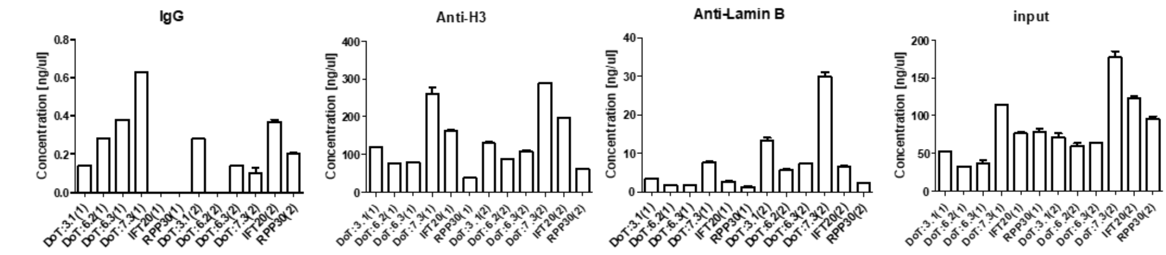


**E**

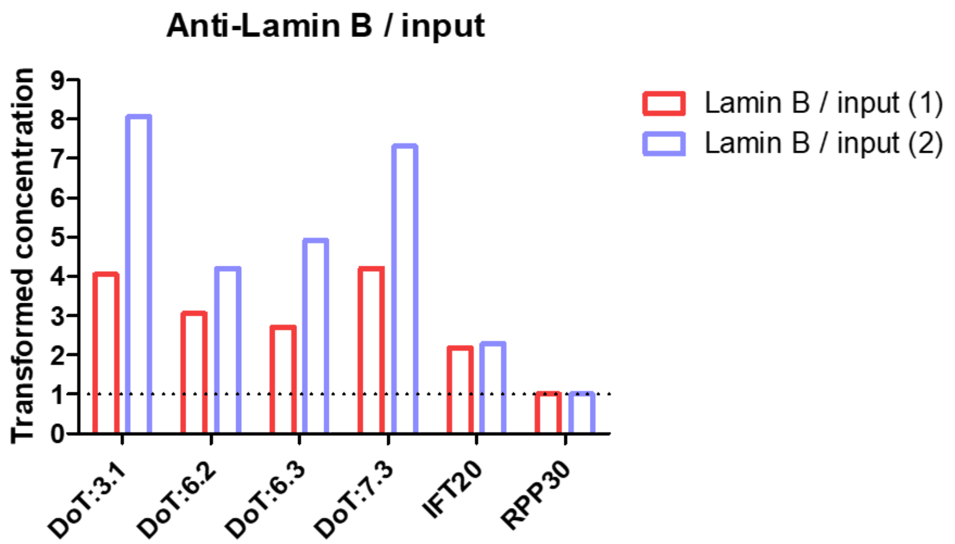

Supplementary Figure S10: Targeted LADs' sites show increased levels of Lamin B. A) Schematic depiction of Chromatin Immunoprecipitation (IP) with ddPCR (ChIP-ddPCR) workflow. After sonication, a part of the lysate was de-crosslinked and loaded on a gel to check the fragment sizes. Then, lysates were divided into input samples and samples for immunoprecipitation. Next, after immunoprecipitation of IP samples,  IP and input samples underwent de-crosslink and purification on the columns, and were analyzed by ddPCR. B) Sonication check. Sonicated samples were after de-crosslink and purification on columns loaded on a 1% agarose gel. Every sample is loaded twice with different loading dye (a white cloud on a gel) to assess the highest intensity of fragments. For the next manipulations, the samples of the same sonication were pooled. C) Digital Droplet PCR concentrations [ng/μl] in the tab. D) Concentrations of Lamin B, input, and H3, respectively, obtained by Digital Droplet PCR. E) Transformed levels of Lamin B related to the inputs. Chromatin fragments' sizes prior to immunoprecipitation are critical for the values. The shorter fragments (sonication #1), which are more target-site specific, exhibit smaller differences than the longer fragments (sonication #2), which are more domain-specific.

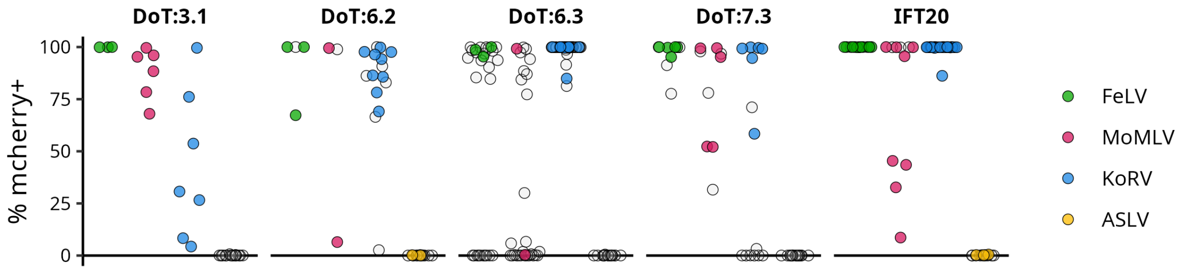


Supplementary Figure S11: Clones tested for targeted knock-in. Clones without verified targeted knock-in are represented by empty circles.

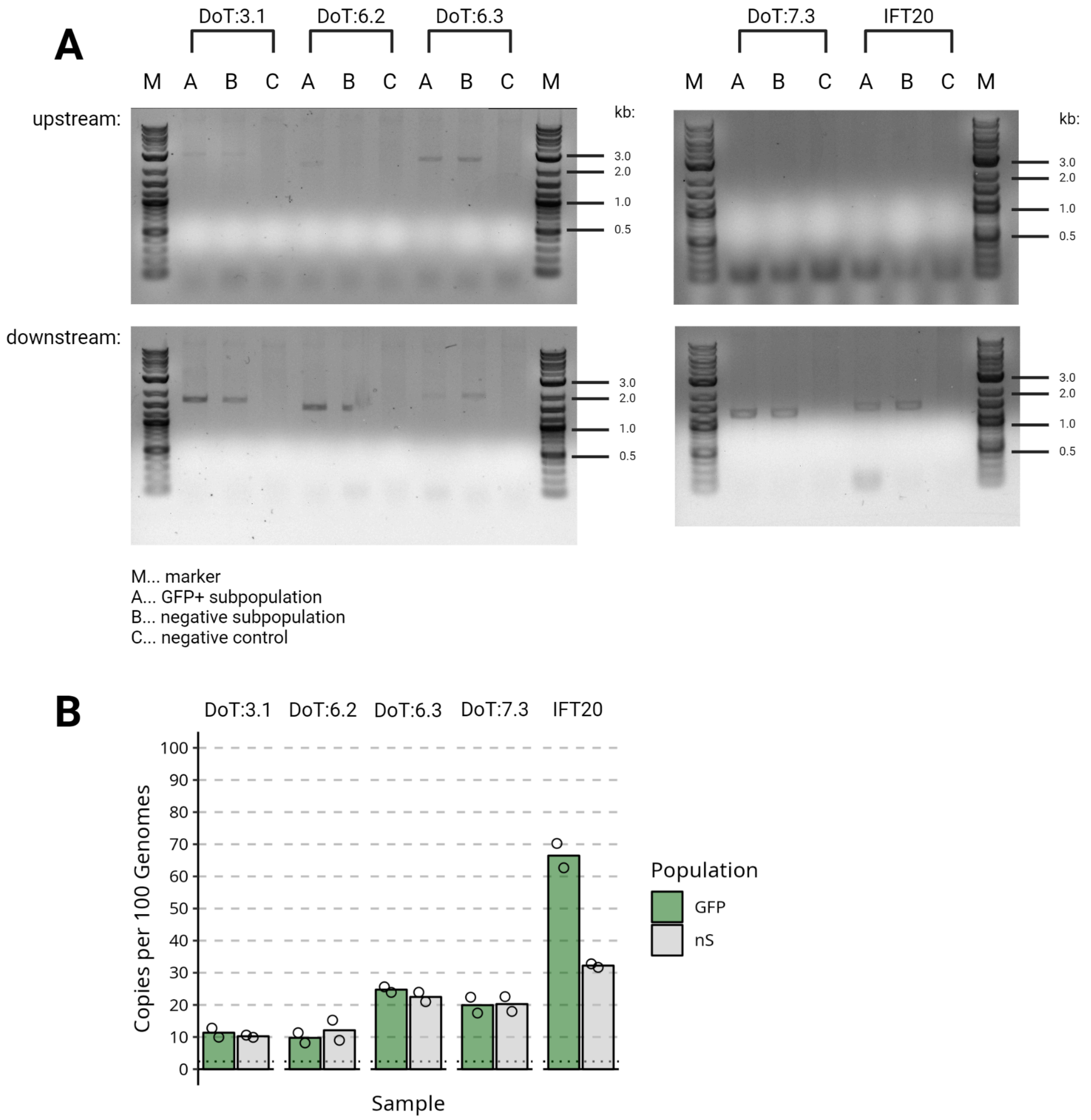


Supplementary Figure S12: ASLV Targeted Knock-in in Negative Populations. A) We amplified upstream and downstream genome-insert junctions after cotransfection CRISPR-Cas9 and homologous-arms-flanked ASLV mini-genome vectors in GFP+ and GFP- subpopulations. B) Analysis of ASLV-vector copy number in GFP+ population and subsequent mCherry- population by digital droplet PCR.

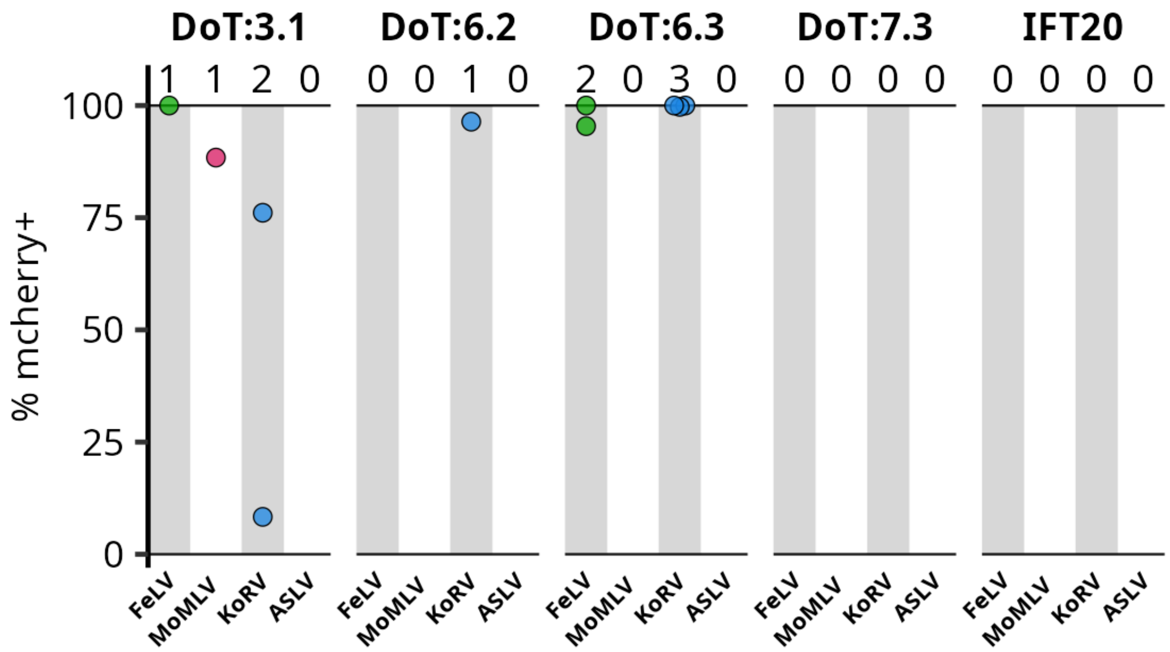


Supplementary Figure S13: mCherry expression in clones with double knock-in in target sites.

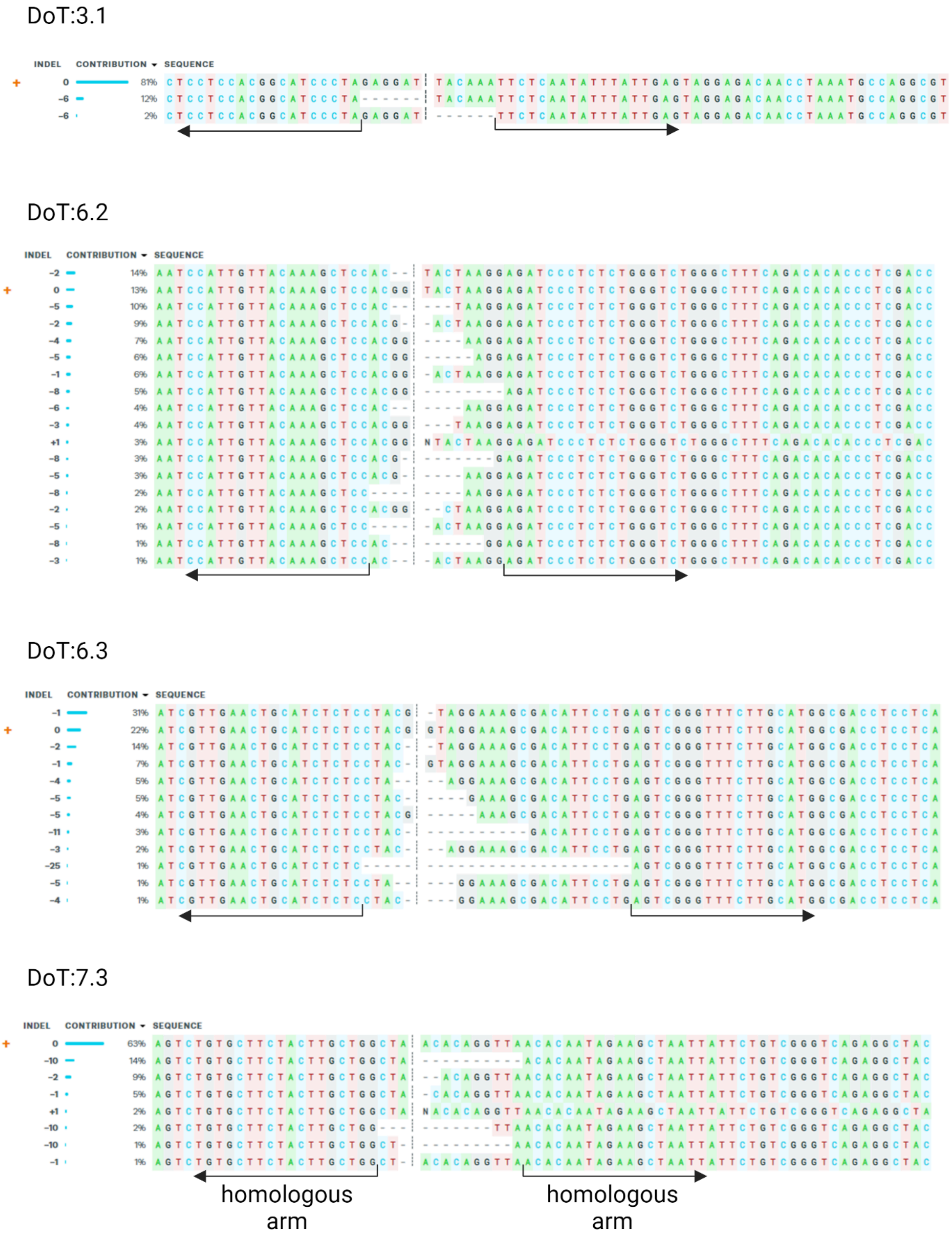


Supplementary Figure S14: LAD target sites' CRISPR cutting efficiency. Apart from assessing the efficiencies, the cutting profiles were used for homologous arms design. Arrows indicate the beginning of homologous arms adjacent to proviruses.
